# Generation of ordered protein assemblies using rigid three-body fusion

**DOI:** 10.1101/2020.07.18.210294

**Authors:** Ivan Vulovic, Qing Yao, Young-Jun Park, Alexis Courbet, Andrew Norris, Florian Busch, Aniruddha Sahasrabuddhe, Hannes Merten, Danny D. Sahtoe, George Ueda, Jorge A. Fallas, Sara J. Weaver, Yang Hsia, Robert A. Langan, Andreas Plückthun, Vicki H. Wysocki, David Veesler, Grant J. Jensen, David Baker

## Abstract

Protein nanomaterial design is an emerging discipline with applications in medicine and beyond. A longstanding design approach uses genetic fusion to join protein homo-oligomer subunits via α-helical linkers to form more complex symmetric assemblies, but this method is hampered by linker flexibility and a dearth of geometric solutions. Here, we describe a general computational method that performs rigid three-body fusion of homo-oligomer and spacer building blocks to generate user-defined architectures, while at the same time significantly increasing the number of geometric solutions over typical symmetric fusion. The fusion junctions are then optimized using Rosetta to minimize flexibility. We apply this method to design and test 92 dihedral symmetric protein assemblies from a set of designed homo-dimers and repeat protein building blocks. Experimental validation by native mass spectrometry, small angle X-ray scattering, and negative-stain single-particle electron microscopy confirms the assembly states for 11 designs. Most of these assemblies are constructed from DARPins (designed ankyrin repeat proteins), anchored on one end by α-helical fusion and on the other by a designed homo-dimer interface, and we explored their use for cryo-EM structure determination by incorporating DARPin variants selected to bind targets of interest. Although the target resolution was limited by preferred orientation effects, small scaffold size, and the low-order symmetry of these dihedral scaffolds, we found that the dual anchoring strategy reduced the flexibility of the target-DARPIN complex with respect to the overall assembly, suggesting that multipoint anchoring of binding domains could contribute to cryo-EM structure determination of small proteins.

## Introduction

There has been considerable interest in designing novel protein assemblies, for example to develop cryo-electron microscopy (cryo-EM) scaffolds to aid in structure determination^1,2^ and protein nanoparticles with antigen display capabilities as vaccine candidates^3,4^. The symmetric assembly design paradigm uses symmetry and either genetic fusion or a designed interface to fix the orientation of symmetric homo-oligomeric building blocks within the overall assembly. The genetic fusion approach was originally demonstrated with the creation of a tetrahedral protein nanocage and fiber^5^ and has since been used to generate two-dimensional layers^6^, a porous cube^7^, additional tetrahedra^8,9^, octahedra^10^, and icosahedra^11,12^. The fusion procedure is relatively straightforward, not inherently destabilizing, and has perfect specificity as the interaction partners are fused. Despite its success and relative simplicity, several aspects of the genetic fusion approach have limited its utility compared to methods that employ non-covalent protein-protein interface design^4,13-17,^. Interface design produces vastly more geometric solutions than genetic fusion, because the available alignment geometries for fusion at helical termini are spatially discrete and finite in number. This reduces the number of possible structures accessible by fusion and increases the difficulty of building into an assembly any particular building block of interest for a given application. In contrast, the adjustable degrees of freedom (rotation and translation) accessible through noncovalent interface design have continuous ranges^18^, so the set of valid geometric solutions is technically unlimited. Other issues with the fusion approach are that the termini must be accessible and that flexibility is often introduced at the point of fusion, even with α-helical linkers and certainly with disordered linkers. In the best cases, model-deviations are subtle^8,19^, however varying levels of unintended assembly products are also commonly observed.

Genetic fusion has been applied to the creation of cryo-EM scaffolds^1,2,20-22^: if a small target protein can be immobilized and rigidly bound onto a larger symmetric assembly, EM particle images can be more readily aligned and classified than those of the target protein alone. Structures that would normally be too small to analyze would then become amenable to structure determination. Yeates and colleagues demonstrated the potential of this approach by fusing a DARPin^23,24^ to the outside of a previously designed protein nanocage^1^; the bound GFP target was resolved at 3.8 Å resolution with only a single α-helical fusion anchoring the DARPin.

## Results

### A computational method for rigid three-body / multi-domain symmetric fusion

We set out to develop a computational method for generating symmetric assemblies by gene fusion that explores vastly more combinations than previous methods and enforces rigid connections between the building blocks. Previous genetic fusion studies have focused on fusing symmetric oligomeric building blocks together at their N and C termini. Furthermore, it has been shown that the rigidity of DARPin fusions depends on the connecting helix being shared, i.e. being part of both domains being connected^25,26^. We reasoned that a much larger set of possible configurations could be generated by (1) incorporating a variable length rigid monomeric protein spacer between the two oligomers, and (2) allowing fusions at internal residues (not just the termini). The number of accessible configurations increases from on the order of N(oligomer_1_) * N(oligomer_2_) for direct fusion at building block termini, to N(oligomer_1_) * N(oligomer_2_) * N(spacer) with the addition of spacer domains, to on the order of N(oligomer_1_) * N(fusion sites per oligomer_1_) * N(oligomer_2_) * N(fusion sites per oligomer_2_) * N(spacer) * N(fusion sites per spacer) * N(fusion sites per spacer) when internal fusion sites are allowed, a very considerable increase. To ensure rigid structurally coherent junctions between the building blocks, we only allow fusion via alignment and superposition of shared helices from both building blocks being fused and disallow α-helical extension. Most globular proteins are destabilized by truncation in the midst of secondary structure elements; to maintain stability, we use idealized repeat protein building block spacers (and oligomers, when possible), where every repeat unit is identical – such proteins are amenable to truncation or fragmentation without undermining folding and stability^27-30^.

Geometric matches to the desired symmetry are identified for each (oligomer_1_, spacer, oligomer_2_) tuple by the following procedure, which is illustrated schematically in Figure 1. First, all rigid body transforms (T_1_) on oligomer_1_ are enumerated that superimpose one of its helical segments onto one from the spacer. Rigid body transforms T_2_ (applied to oligomer_2_) are calculated identically, to identify possible fusions between oligomer_2_ and the spacer. Second, for each (T_1_, T_2_) instance combination, the transforms are applied to both oligomers and the arrangement of the axes of the two oligomers are tested for compatibility with the target architecture. For D_2_ symmetry, they must intersect at 90 degrees or for D_3_, 60 degrees. Next, the symmetry is idealized by rotating each homo-oligomer about its corresponding shared-helix-segment center-of-mass so that the axes meet perfectly at the required angle. In cases where the applied rotation (a measure of non-ideality) is less than the configured error tolerance, 3D models of the resulting assemblies are built by superposition of the repositioned structures. Models are discarded in which the fusions truncate homo-oligomer interface residues, as are models with backbone clashes; sidechain clashes are acceptable, since redesign can eliminate them.

**Figure 1.**
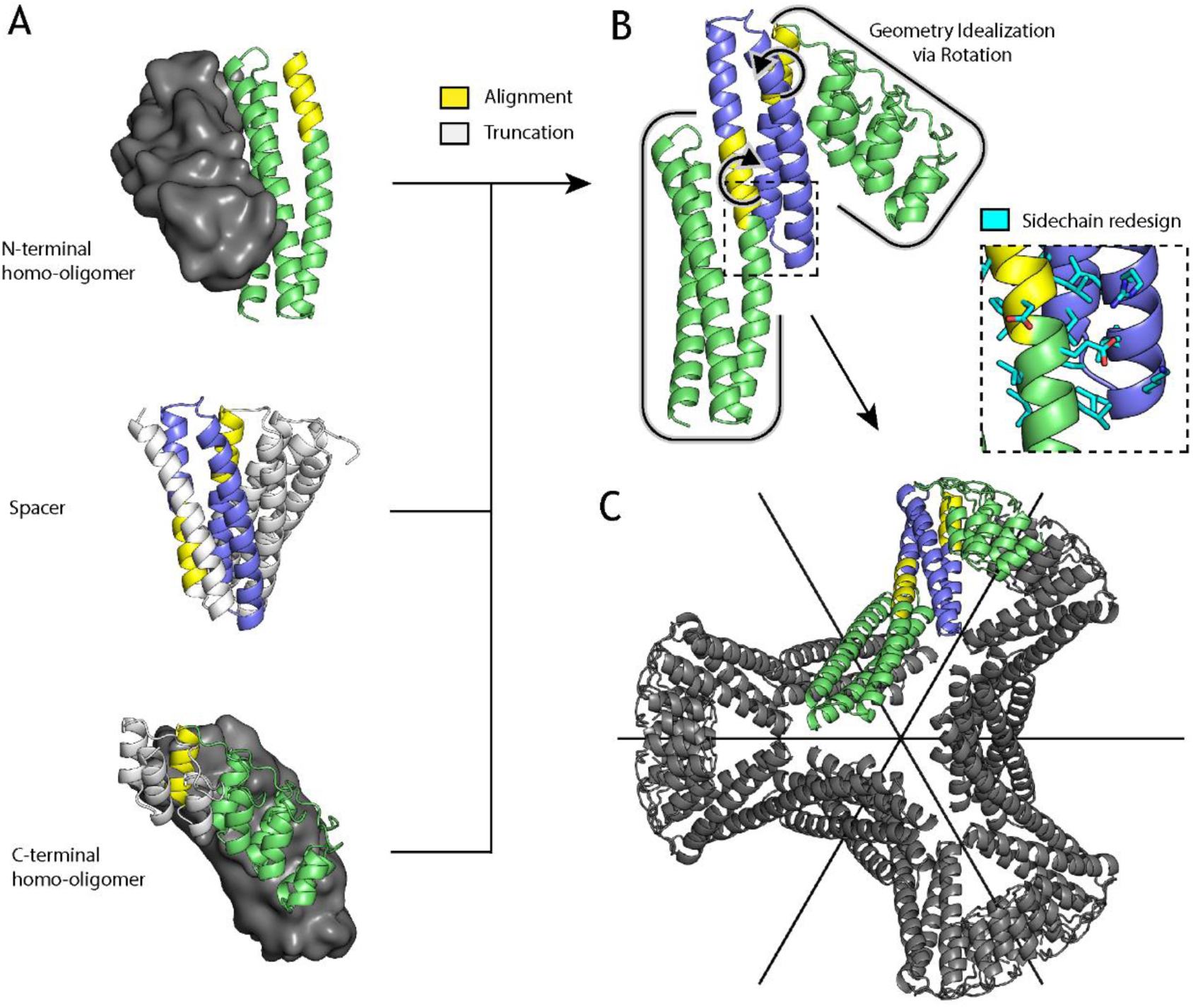
Illustration of the tripartite design strategy for the D_3_ architecture. The final structure is composed of (A) two homo-oligomers (dimers, top and bottom; the partner subunit is shown as a surface in grey) and a de-novo helical repeat protein (middle). All possible non-clashing backbone alignments are geometrically analyzed and filtered to generate (B) a three-component fusion, which is idealized to the target geometry by small rigid-body rotations and redesigned to improve core packing and remove exposed hydrophobics. The result (C) is a D_3_ assembly, with symmetric C_2_ axes (black) that correspond to those of the original homo-dimers and a new C_3_ axis orthogonal and through the center.

In selecting helical segment pairs for fusion in the first step of the above procedure, we use several criteria to try to ensure fusion rigidity. First, we require superpositions with a low backbone R.M.S.D. (default 0.5 Å) over multiple residues (default 8 minimum) to ensure the shared helix geometry is consistent with both building blocks. The stringency is controlled by user-configurable backbone R.M.S.D. and overlap-length parameters. Unlike traditional symmetric fusion with α-helical linkers, overlapping fusion preserves proximity and sidechain packing of the fusion region with remaining secondary structures from the original building blocks, which reinforces the structure. For the sake of efficiency, the R.M.S.D. and overlap-length thresholding occurs early in the procedure during geometric-match identification, when alignments are initially computed. Second, we rank fusions according to a rigidity metric, which counts the number of putative sidechain contacts between secondary structures of previously separate building blocks. This ensures that significant cross-building-block contacts can be made to buttress the interactions along the shared helix. This step also occurs prior to sidechain redesign, so Cα-Cβ vectors are used in place of contact counts as a measure of designability. Third, we redesign with Rosetta the regions adjacent to the new building block junctions to eliminate steric clashes and improve packing between newly joined regions. Fusion-related truncation often exposes hydrophobic residues that were previously buried, so these regions must be redesigned as well.

### Design and characterization of D_2_ and D_3_ symmetric oligomers

As a proof of concept, we applied the rigid three-body fusion method to design dihedral symmetric assemblies from de novo designed repeat protein monomers and oligomers. Two different designed C_2_ dimers were fused as described above with a repeat protein monomeric spacer such that the C_2_ axes intersect at 90 degrees for D2 structures or 60 degrees for D3 structures (Figure 2). Two different design rounds were performed targeting D_2_ and D_3_ symmetries. Even in the second design round, where fewer building blocks and additional selection criteria were imposed, millions of tripartite alignment combinations were scanned per symmetry, nearly ten thousand fusions matched the target geometry within the angular error tolerance (5 degrees), and several hundred passed junction rigidity metrics. The sequences at the junction regions in these assemblies were then optimized using Rosetta, as described in the Methods. In brief, the full symmetric assemblies were generated from the single-chain asymmetric unit and a single round of sidechain redesign with the beta_nov16 score-function was performed on the junction residues, while disallowing the introduction of cysteines, prolines, or methionines.

**Figure 2.**
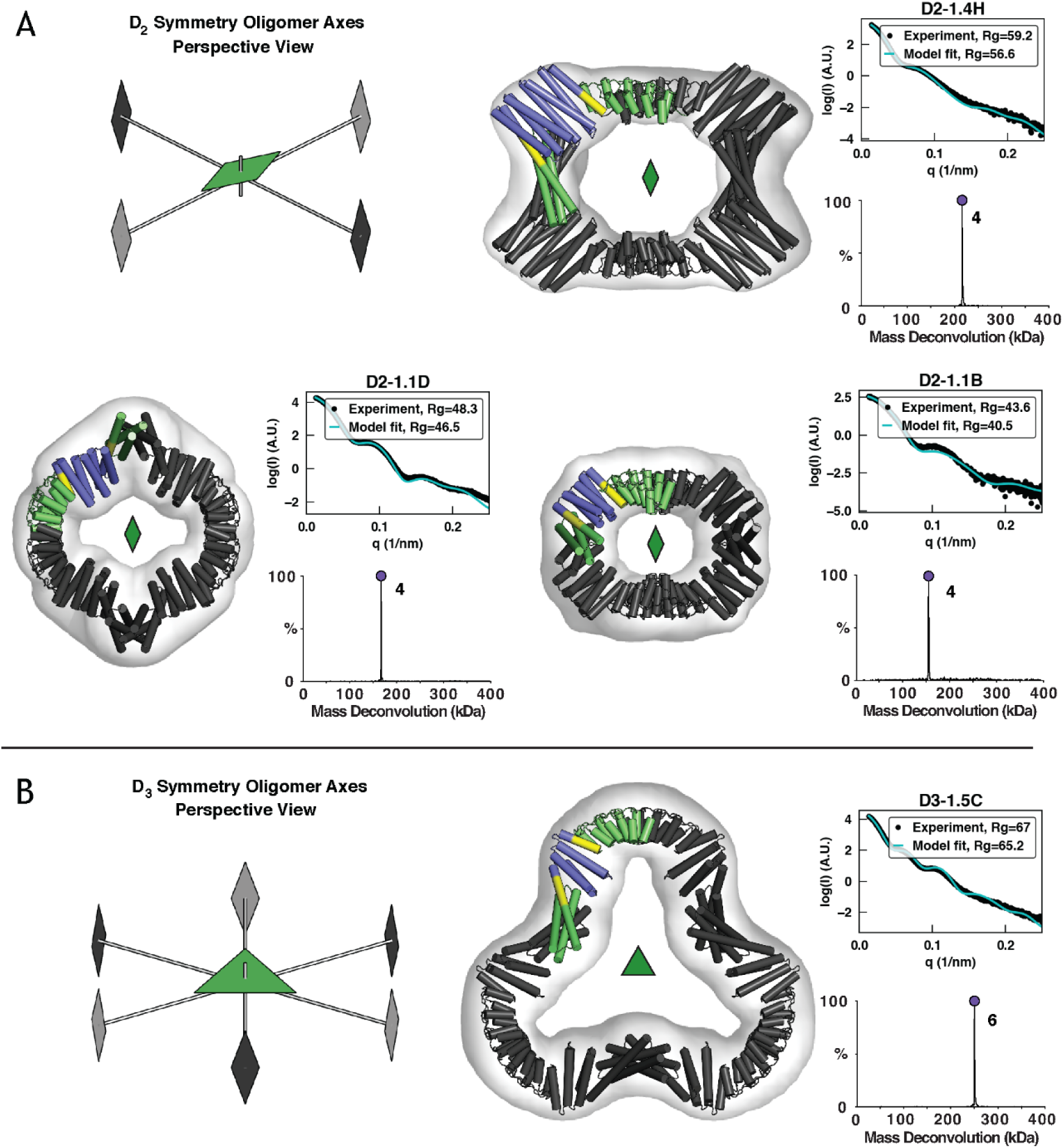
Characterization of first round designs. EM, native-MS, and SAXS experiments are consistent with the formation of intended architectures for four designs: (A) three D_2_ designs and (B) one D_3_ design. Negative-stain 3D reconstructions are overlaid by design models, whose asymmetric unit is colored according to its constituent building blocks (N and C-terminal oligomers, green; DHR, blue; shared alignment, yellow). Native-MS deconvolutions show the relative abundance of the determined masses and the peaks are labeled with their assigned oligomeric states. SAXS plots compare the theoretical (cyan) and experimental (black) scattering intensities (log scale) as a function of q, as well as radius of gyration (Rg), in Ångströms.

In a first design round, 28 D_2_ and 9 D_3_ assemblies which matched the geometric selection criteria and had low energy junctions were selected for experimental characterization and recombinantly produced in *E. coli*. Of these 37 designs, seven (three D_2_ and four D_3_) were both soluble and eluted as single monodisperse peaks by size exclusion chromatography (SEC) (Figure S1). Scattering profiles and radius of gyration (Rg) determined from solution X-ray scattering (SAXS) data were consistent with design models for the three D_2_ designs and for two of the four D_3_ designs (Figure 2 and S2). Of these designs, native mass spectrometry (native-MS) verified the expected oligomeric states for all of the assemblies. Despite the limited dataset, clear trends emerged: for example, all five designs corroborated by SAXS data incorporated a C-terminal ankyrin homo-dimer, which was present in only 60% of the tested designs. In addition, four of five SAXS and native-MS corroborated designs incorporated the same N-terminal three-helix homo-dimer “rop20”, despite rop20 only being present in ∼40% of tested designs. The set of 12 designs that combined both an N-terminal rop20 and C-terminal ankyrin dimer contained four of the five successes. Meanwhile, no designs using hairpin helical bundle dimers were successful.

A second round of design was performed with the same procedure, this time using mostly the rop20 helical bundle at the N terminus and any of the three similar ankyrin dimers at the C-terminus that had been successful in the first round. We also introduced two additional design constraints with the aim of improving the designs’ suitability as cryo-EM DARPin scaffolds, whose use is discussed in the next sections. First, structures with fewer total secondary structures (helix count) were selected, anticipating that they would be more rigid. Although the design method reinforces the point of fusion, the de-novo helical repeat (DHR) spacers and ankyrin dimer building blocks have small cross-sectional areas, so each additional repeat likely adds flexibility. Second, we selected designs with the ankyrin binding groove facing away from the assembly center (Figure S6), to reduce the chance of steric hindrance in multivalent target binding. 31 D_2_ and 24 D_3_ designs were ordered, of which 15 and 12, respectively, had high levels of soluble expression and single major peaks by SEC. SAXS and native-MS agreed with the design models of four D_2_ designs and four D_3_ designs (designs D2-21.29 and D2-21.30 have 85% sequence identity and were not considered independent successes; only D2-21.29 was fully characterized) (Figure 2).

With the exception of design D2-21.22 and D3-1.5A2 (Figure S3), which may exhibit lower stability, negative-stain EM 2D class averages and 3D reconstructions recapitulated the expected shape for all designs that passed both native-MS and SAXS screening. Most designs have a pronounced central cavity that makes their top views readily identifiable in micrographs. An unusual structural aspect of our D_3_ designs is their subunit connectivity; in natural D_3_ architectures, subunits related through C_3_ rotations normally make direct contact. In almost all of our designs, they do not – the C_3_ axis is an emergent property of the assembly. Another notable feature of our designs is that they were produced in large part from building blocks without crystal structures. Of the ten successful and sequence-independent designs, two use not a single crystal-verified building block (D2-1.4H and D3-19.24) and another seven use only one crystal-verified building block (Table S2).

### Cryo-EM of coassembled DARPin and GFP

In prior studies, imaging scaffolds have been constructed through fusion of a DARPin onto existing symmetric protein assemblies via a shared helix or α-helical extension^1,2^. Our constructs differ in that they incorporate a designed ankyrin homo-dimer in addition to fusion, so the ankyrin (or DARPin post interface-grafting) is held in place by two separate mechanisms (back-to-back dimerization and lateral fusion) that should both contribute to rigid placement. To test our scaffolds, we grafted the surface binding residues from a GFP-binding DARPin^24^ into six constructs: D2-1.1D, D2-1.4H, D3-1.5C, D2-21.8, D2-21.29, and D3-19.20, taking care not to alter the ankyrin homo-dimerization interface in the base construct (Figure S2). As ankyrins are repeat proteins, the DARPin alignment can be shifted up or down by adding or removing one or more consensus repeats, which we did to create variants. After SEC purification and cryo-EM of GFP-scaffold co-complexes, a majority of constructs exhibited high levels of preferred orientation or outright aggregation (Table S3), but D2-1.4H and D2-21.8 derivatives were promising. D2-1.4H.GFP.v1 is derived from the first round design scaffold D2-1.4H and was reconstructed at 4.3 Å resolution for the full complex and GFP target separately (Figure 4) – just as well as the core scaffold (Figure S7A). Second-round design derivative D2-21.8.GFP.v2 was resolved at lower resolution of only 6.0-7.0 Å overall for the co-complex with the GFP target (Figure S7B).

**Figure 3.**
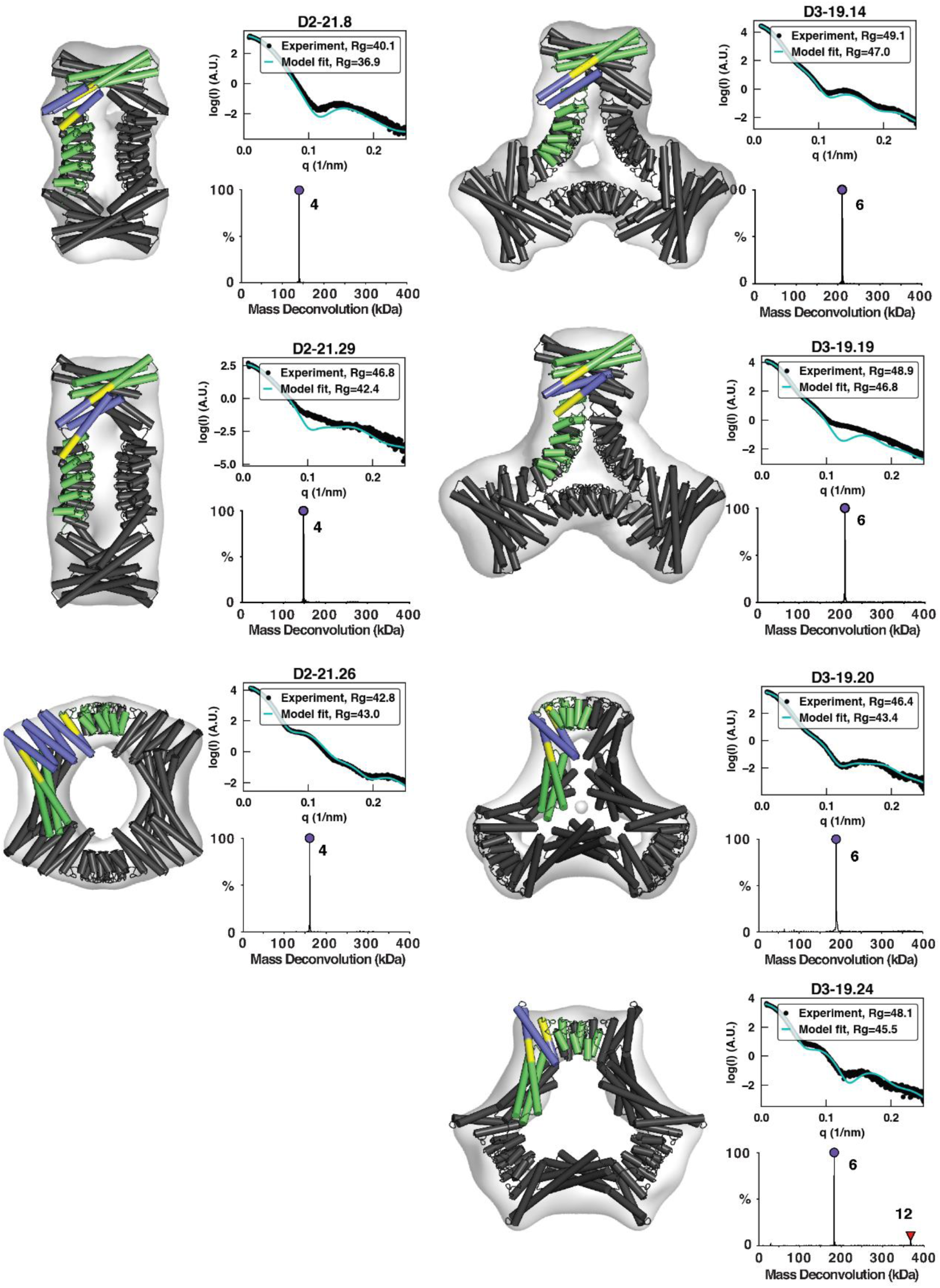
Characterization of second round designs. EM, native-MS, and SAXS experiments are consistent with the formation of intended architectures for six designs. Negative-stain 3D reconstructions are overlaid by design models, whose asymmetric unit is colored according to its constituent building blocks (N and C-terminal oligomers, green; DHR, blue; shared alignment, yellow). Native-MS deconvolutions show the relative abundance of the determined masses and the peaks are labeled with their assigned oligomeric states. SAXS plots compare the theoretical (cyan) and experimental (black) scattering intensities (log scale) as a function of q, as well as radius of gyration (Rg). Native-MS for D3-19.24 shows a small amount of 12-mer, likely formed through association of two designed hexamers.

**Figure 4.**
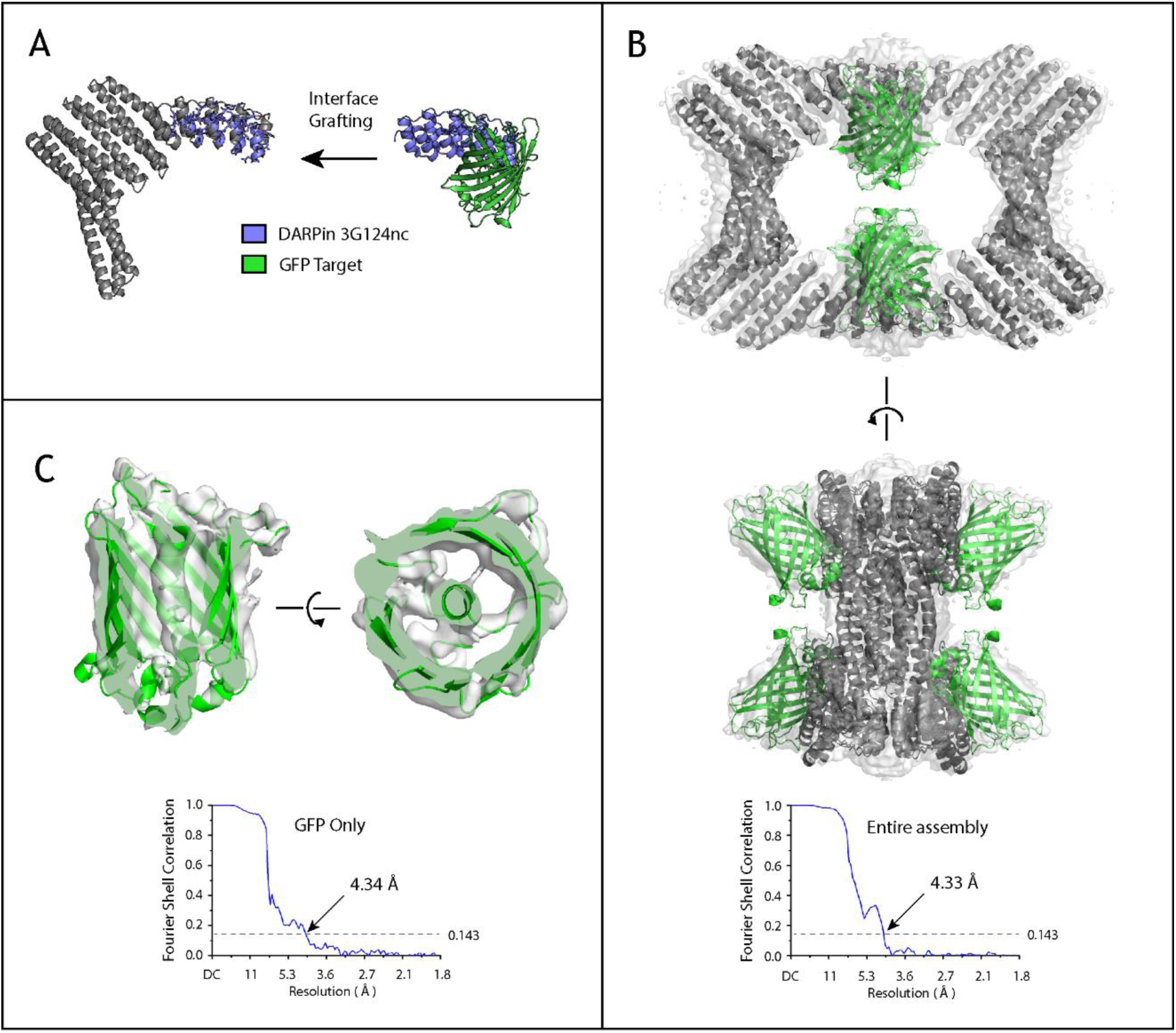
Cryo-EM characterization of DARPin-grafted scaffold with GFP. **(A)** Surface residues of the GFP-binding DARPin “3G124nc” ^24^ are grafted onto the assembly subunit of D2-1.4H, while preserving core residues and homo-oligomer interfaces, to form a hybrid (D2-1.4H.GFP.v1)that retains self-assembly and also binds GFP. **(B)** Cryo-EM density of the co-assembled complex is inlaid with cartoon models. The corrected Fourier shell correlation (FSC) curves are calculated by the program cryoSPARC, shown in blue. Using the standard FSC=0.143 criterion, the nominal resolution is 4.33 Å. **(C)** GFP density with the core scaffold masked. The GFP portion has nominal resolution 4.34 Å.

### Cryo-EM of coassembled DARPin-scaffold with Human Serum Albumin (HSA)

All cryo-EM scaffolding studies based on the GFP-binding DARPin had the benefit of an available co-complex crystal structure, but in the general use case co-complex structural information will be absent (the purpose of the scaffold being to facilitate structure determination) and we sought to assess the feasibility of integrating DARPins in such a scenario. Towards this aim we incorporated an unpublished DARPin sequence targeting human serum albumin (anti-HSA DARPin “C9”) into second-round scaffolds D2-21.8 and D2-21.29. As before, the DARPin sequence was aligned and grafted onto the assembly scaffold, taking care to only graft surface residues and not to mutate the ankyrin homo-dimer interface. DARPin residues in the hydrophobic core that differed from those in the scaffold were not grafted to avoid disrupting the homo-dimer interface. Four sequence-grafted designs were expressed (2 scaffolds x 2 variants where the grafted surface residues are shifted by a repeating unit of the ankyrin). One design based on the D2-21.8 base scaffold and the second repeat-unit-shifted variant proved more soluble than the others (D2-21.8.HSA-C9.v2) and SEC with SDS-PAGE confirmed binding to HSA (which appeared to be sub-stoichiometric based on SDS-PAGE of column fractions).

The HSA complex was successfully reconstructed by cryo-EM to 5.5 Å resolution (Figure 5). In line with observations on stoichiometry, the HSA-binding mode sterically precludes full binding site occupancy and each face of the dihedral ring has one HSA instead of two. This results in two predominant species where two of four binding sites are occupied, either across from one another or along the diagonal – image classification and refinement focused on the former. The core scaffold map density is greater than that of the HSA density, so for the cutoff level shown, each HSA model can be fit fully within the map, but the scaffold core density appears overly large; this effect is exacerbated by preferred orientation. Still, the placement and DARPin binding site is clear. Aromatic residues on the DARPin overlap a hydrophobic patch in domain II of HSA, away from the site where HSA binds to FcRn, whose binding enables HSA escape from endosomal degradation and long serum half-life. This non-interference indicates that the HSA DARPin “C9” may serve as a fusion domain for half-life extension. The process also demonstrates the feasibility of integrating DARPins for determining interaction sites with the target, even with the added complexity of maintaining the original homo-dimer interface.

**Figure 5.**
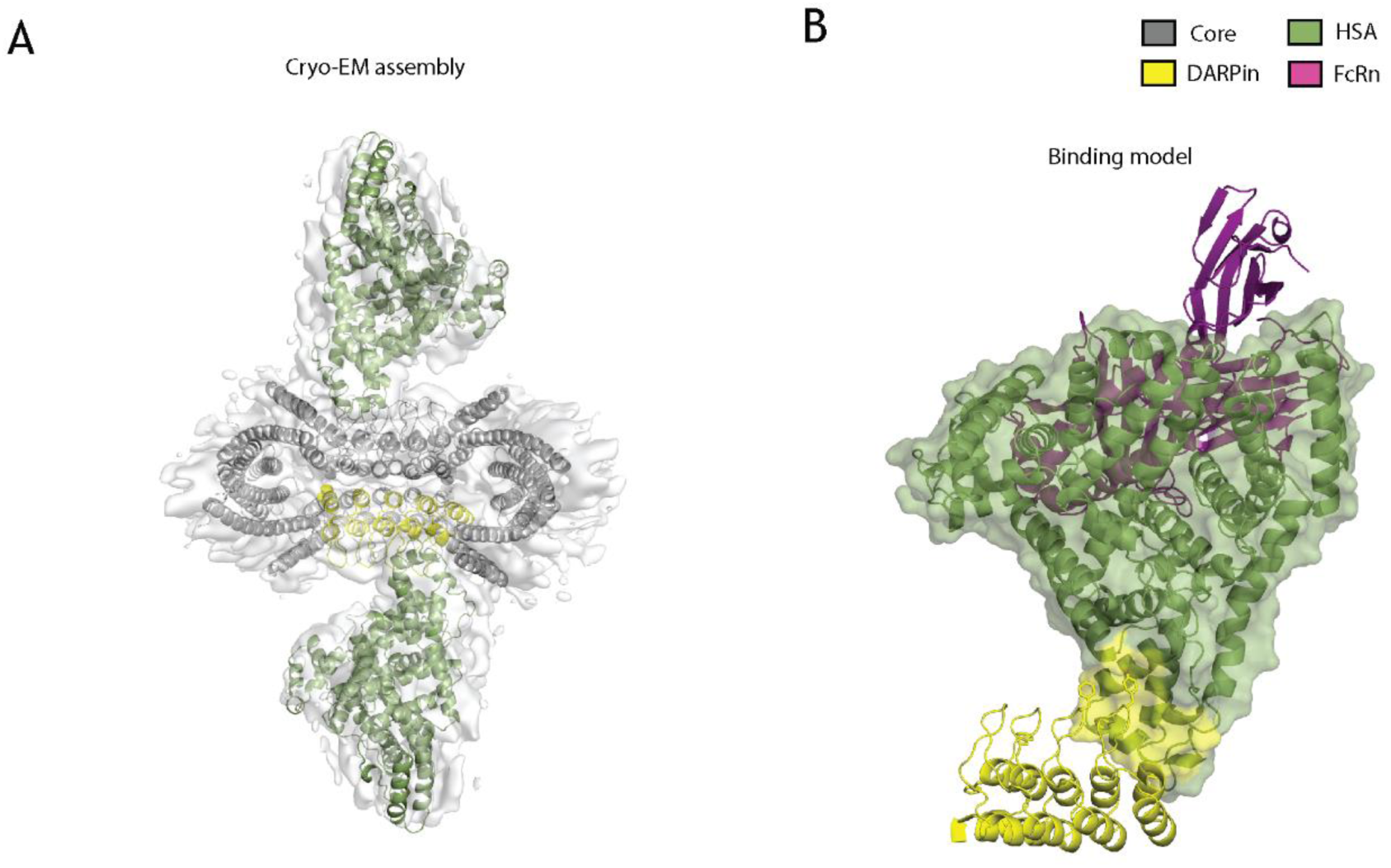
Characterization of anti-HSA DARPin assembly in complex with HSA. A) Cryo-EM of scaffold 21.8.HSA-C9.v2 and HSA co-complex with embedded scaffold design model and HSA crystal structure (PDB 1BJ5). A single HSA-binding DARPin (of four total and two that bind per co-complex) is highlighted in yellow. B) A model of the relative HSA-binding positions of the DARPin and FcRn built by superposition of this cryo-EM structure (DARPin scaffold + HSA) and an existing crystal structure (HSA + FcRn, PDB 4K71). The HSA surface representation is colored within 10 Å of the DARPin.

## Discussion

The new multi-fusion method introduced here specifically remedies two of the long-standing drawbacks to fusion-based assembly construction: the low number of geometric solutions and linker flexibility. Our results demonstrate the feasibility of performing multiple fusions, ranking, and redesign in a single pass to produce intended architectures with low levels of off-pathway assembly. It remains to be seen how well the method performs for more complex and multi-component assembly architectures. The role of rigidity is also an open question: the cryo-EM application would clearly benefit from a more rigid overall structure, but the design method may require a measure of component flexibility to compensate for design inaccuracies around the redesigned fusion. It is notable that all successful designs in the present work incorporated at least one, usually two, and occasionally even all three design models lacking high resolution structural validation (SAXS validation only); it is possible that using x-ray crystal structures of the building blocks rather than computational design models might lead to greater success. On the other hand, the success using design models bodes well for future applications of this approach as far more plausible building block structures can be designed than can be solved by x-ray crystallography.

We had hypothesized that DARPins would be more rigidly embedded in our new designed assemblies than in the previous designs by Liu et al. ^1,22^, as in the present work they form structural components of the assembly and are tethered by either a fusion or designed interface at either end. However, the resolution we achieved was half an Ångström lower. In the previous study, the core scaffold was resolved to 2.9 Å and the GFP target was resolved to 3.8 Å, which indicates some flexibility of the DARPin position relative to the core scaffold. In the present study, the GFP target was resolved at 4.3 Å resolution, just as well as the core scaffold. Hence, the secondary stabilization provided by the homo-dimer interface appears to have been effective in rigidifying the DARPin placement with respect to the rest of the scaffold, but this was counteracted by some combination of inherent flexibility in the core dihedral scaffold and preferred orientation in cryo-EM, which limited the achieved resolution. We expected that the shorter closure path selected for in the second round designs would improve rigidity and produce better scaffolds than those from the first round of design. Instead, the second round designs showed a greater tendency toward aggregation and preferred orientation in cryo-EM (Table S3), which may be rooted in our selection criteria; while the shorter closure paths may indeed increase rigidity, it also biased toward smaller designs that are likely more sensitive to destabilization when grafting in the hydrophobic DARPin interface. In line with this hypothesis, D2-1.4H.GFP.v1 yielded the best results and also contains the largest (in terms of cross-section width and total mass) DHR building block among designs tested by cryo-EM. While the scaffolds here fall short of recapitulating or improving upon the 3.8 Å GFP resolution achieved by Liu et al., the use of secondary stabilization through interface design could still yield improved results for cryo-EM if used in conjunction with bulkier building blocks and higher order symmetry. Beyond scaffolds for cryo-EM structure determination, the rigid three-body fusion method demonstrated here provides a general strategy for producing arbitrary geometries and facilitates exploration of the protein nanomaterial design space.

## Materials and Methods

### Computational design

A custom software library was built with the .NET Framework, which includes functionality for PDB parsing, alignment, symmetry/patterning, clash and contact checking, structure editing, and running the multi-domain fusion algorithm. The several parameters that control fusion were assigned based only on manual curation of outputs during testing in-silico and are likely not optimal for all scenarios. In particular, an 8-residue minimum overlap-length was selected because the idealized ankyrins used in this study have short helices compared to those of DHRs and helical bundles, but longer overlaps might be desirable with other starting components. Likewise, a lower angular-error tolerance might increase the success rate of tested designs, but it was kept at a moderately high 5 degrees, because the lack of crystal structures for so many designs introduced uncertainty about the initial model accuracy, so a tight angle tolerance would have been somewhat arbitrary.

As the method creates a much larger solution space than direct fusion, optimizations were necessary to keep runtimes reasonable while still exhaustively enumerating geometries. The most impactful optimization eliminates redundant alignments by greedily expanding the alignment windows of any valid 8-residue alignment until the R.M.S.D. threshold is exceeded or either secondary structure element ends – all shorter alignment windows contained within the expanded alignment need not be examined. The result is that fewer alignment combinations are considered than if every 8-residue window were examined and nearly identical outputs are largely avoided. The protocol could be run on all de-novo building-block combinations for a target geometry in less than 24 hours on a quad-core laptop; higher parallelism would produce a speedup accordingly.

The fusion output models were redesigned by Rosetta with a simple RosettaScripts protocol (Text File S1), involving only two Movers (operators that modify a design model): SetupForSymmetry and SymPackRotamersMover. These Movers respectively recreate the full symmetric assembly from the input single-chain asymmetric unit and redesign those residue side chains that were identified by output files in the Resfile format. After the initial sidechain redesign pass, models deemed promising by a combination of total score and manual inspection were subjected to one or more additional redesign passes with the same protocol, but with user-generated Resfiles, to eliminate exposed hydrophobic residues, revert residues to their original wildtype identity, or mutate Rosetta-designed glycines to alanines within helices to improve helical propensity. The beta_nov16 score function was used throughout.

The input structure set consisted of 20 homo-dimer and 42 DHR spacer proteins already verified within the lab, with 5 homo-dimers and 15 DHRs having been previously published with solved crystal structures available in the Protein Data Bank^18,31,32^, (Table S4). Two designed crystal structures were unintentionally omitted from the input set (2L4HC2_4 and 3L6HC2_2 from Boyken et al). Two-helix dimers were removed from the scaffold set in the second round of design, because better results were obtained from three-helix dimers. The third helix leads to a larger hydrophobic core than exists in the two-helix dimers, which we expect leads to a higher degree of degree of order even in the monomeric form and might help to avoid aggregation and misassembly. The other type of successfully incorporated dimer was based on ankyrins. Although very similar, the minor binding orientation differences between the three ankyrin homo-dimers was sufficient to make all three useful in finding distinct geometric solutions.

### Cryo-EM of coassembled DARPin and GFP

Electron microscopy grids were prepared at 4°C at 100% humidity using vitrobot (FEI). In brief, 3 µl of purified sample at 1.0 mg/ml was applied to glow-discharged Quantifoil 200 mesh R1.2/1.3 grid, and was manually blotted with a filter paper (Whatman No. 4) for approximately 3 seconds before plunging into liquid ethane. The grids were screened on a Talos Arctica 200 kV with K3 direct electron detector for ice thickness and sample distribution. Micrographs of the screened grid were collected on a Titan Krios microscope (FEI) operating at 300 kV with energy filter (Gatan) and equipped with K2 Summit direct electron detector (Gatan), using data collection program SerialEM^33^. A nominal magnification of 165,000x was used for data collection, corresponding to a pixel size of 0.865Å at specimen level, with the defocus ranging from −1.0 μm to −3.0 μm. Movies were recorded in superresolution mode, with a total dose of −60 e^−^/Å^2^ and dose rate of 8.4 electron per pixel per second and fractionated into 40 frames. Movies were decompressed and gain-normalized using the program Clip in IMOD. Raw movies were corrected for beam-induced motion and binned by two using MotionCor2^34^, and exposure-filtered in accordance with relevant radiation damage curves^35^. The CTF estimation was performed with GCTF^36^ on non-dose weighted micrographs. Micrographs with high CTF Figure of Merit scores and promising maximum resolution (better than 3.9 Å) were selected for further processing (total 1532 micrographs). Several rounds of autopicking using combinations of different references and manual picking were analyzed to determine optimal settings, and yielded similar results. These particles were subjected to iterative rounds of 2D classification, subset selection of high-quality classes, and re-extraction, yielding 138,348 particles from 1023 micrographs, all in RELION 3.0^37^. The initial model was de novo generated and subsequent 3D heterogenous refinement was performed using cryoSPARC^38^. Particles from the best quality 3D class were selected for further processing. The UCSF PyEM package^39^ was used to convert the cryoSPARC coordinates into RELION. The resulting particles were analyzed by 3D refinement, Bayesian Particle Polishing and CTF Refinement in RELION with C1 or D2 symmetry. All the reconstructions were analyzed using UCSF Chimera^40^. The coordinate model was built by breaking the initial design model into domains and rigidly docking these individual protein structures into the EM map using Chimera. Once the orientation was identified, the model was then fit and adjusted manually in Chimera and Coot^41^. The local resolution and final Fourier shell correlation were calculated using Resmap^42^ and cryoSPARC. The core resolution was calculated using the validation function in cryoSPARC.

### Cryo-EM of coassembled DARPin and HSA

Co-complex of 21.8.HSA-C9.v2 with recombinant human albumin (Albumedix™ Veltis®) was purified by SEC. 3 μL of 1 mg/ml of co-complex was loaded onto a freshly glow-discharged (30 s at 20 mA) 1.2/1.3 UltraFoil grid (300 mesh) prior to plunge freezing using a vitrobot Mark IV (ThermoFisher Scientific) using a blot force of 0 and 6 second blot time at 100% humidity and 25°C. Data were acquired using the an FEI Titan Krios transmission electron microscope operated at 300 kV and equipped with a Gatan K2 Summit direct detector and Gatan Quantum GIF energy filter, operated in zero-loss mode with a slit width of 20 eV. Automated data collection was carried out using Leginon at a nominal magnification of 130,000x with a pixel size of 0.525Å. The dose rate was adjusted to 8 counts/pixel/s, and each movie was acquired in super-resolution mode fractionated in 50 frames of 200 ms. 1,011 micrographs were collected with a defocus range between −1.0 and −3.5 μm. Movie frame alignment, estimation of the microscope contrast-transfer function parameters, particle picking, and extraction were carried out using Warp^43^. Particle images were extracted with a box size of 800 binned to 400 yielding a pixel size of 1.05 Å. Two rounds of reference-free 2D classification were performed using CryoSPARC to select well-defined particle images. The selected particles were subsequently subjected to ab initio 3D reconstructions and 3D refinement using CryoSPARC. CTF refinement was used to refine per-particle defocus values. Particle images were subjected to the Bayesian polishing procedure implemented in RELION 3.0. 3D refinements were carried out using non-uniform refinement along with per-particle defocus refinement in CryoSPARC.

### Native mass spectrometry

Sample purity and oligomeric state was analyzed by online buffer exchange MS^44^ using a Vanquish UHPLC coupled to a Q Exactive Ultra-High Mass Range (UHMR) mass spectrometer (Thermo Fisher Scientific) ^45,46^ modified to allow for surface-induced dissociation (SID) similar to that previously described^47^. With the exception of D3-19.14 (50 μM), 1 μL of 25 μM protein in 25 mM Tris and 150 mM NaCl were injected and online buffer exchanged into 200 mM ammonium acetate, pH 6.8 by a self-packed buffer exchange column (P6 polyacrylamide gel, Bio-Rad Laboratories) at a flow rate of 100 μL per min. A heated electrospray ionization (HESI) source with a spray voltage of 4 kV was used for ionization. Mass spectra were recorded for 1000 – 20000 m/z at 3125 resolution as defined at 400 m/z. The injection time was set to 200 ms. Voltages applied to the transfer optics were optimized to allow for ion transmission while minimizing unintentional ion activation, and a higher-energy collisional dissociation (HCD) of 5 V was applied. Mass spectra were deconvoluted using UniDec V4.2.2^48^. Deconvolution settings included mass sampling every 10 Da, smooth charge states distributions, automatic peak width tool, point smooth width of 1 or 10, and beta of 50 (artifact suppression).

### Protein expression and purification

DNA sequences encoding proteins with 6xHis tags were codon-optimized by Genscript and cloned into pET28b+ or pET29b+ vector under the control of a T7 promoter. Plasmids were transformed into BL21(DE3) *E. coli* and plated on LB agar plates. On different occasions, either 50 ml or 500 ml expression cultures were used. 50 ml expression cultures were directly inoculated from plate colonies and grown for 24 hours in Studier’s autoinduction media^49^ with shaking. Alternatively, 5 ml starter cultures in TB were inoculated and grown for 9-12 hours before transfer to 500 ml autoinduction media for 16-18 hours. All growth media was prepared with 100 μM kanamycin as a selection antibiotic.

Expression cultures were spun down for 10 minutes at 4,000 rcf, resuspended in 40 ml TBS (150 mM NaCl, 25 mM Tris) with Pierce protease inhibitor (Product No. A32963), and lysed by sonication. Lysates were centrifuged at 25,000 rcf for 40 minutes to separate the insoluble fraction. The soluble fraction was purified by affinity chromatography over Ni-NTA Agarose (Qiagen) gravity columns. Eluates were concentrated and fractionated by SEC on a Superdex 200 Increase 10/300 GL.

### Negative-stain EM

PELCO 300 mesh Copper grids with Carbon film (Product 01843-F) were glow-discharged and 3μL of sample in TBS was applied to the grid and blotted immediately. 3 μL 2% uranyl formate stain was applied and blotted immediately, twice, and then allowed to dry. Approximately 50 micrographs per construct were recorded on a Thermo Scientific Talos transmission electron microscope operating at 200kV. The known symmetry (D_2_ or D_3_) was applied during reconstruction, except for designs D3-19.14 and D3-19.19, for which C_1_ symmetry was applied (although the design model is D_3_). 3D reconstructions were generated in either RELION or cisTEM^50^.

### SAXS analysis

SAXS data were collected at the SYBILS Beamline (Advanced Light Source in Berkeley, CA) via their Mail-In SAXS program. KNO_3_ was added to buffer solutions in the range of 2 to 5 mM to minimize radiation-damage induced aggregation. Samples were concentrated in Amicon Ultra 0.5ml centrifugal filters and flow-through was used as the background subtraction buffer. For each sample, the average scattering profile was computed, excluding data in the Guinier region for timepoints after radiation damage became observable. The Scatter software was used for analysis; model and experimental Rg values were determined from their respective Guinier region data. Combined datasets (model-vs-experiment) were generated with the FOXS web server^51,52^ for plotting.

## Supporting information

fusion_supplement

## Author Contributions

I.V., Q.Y., S.J.W., G.J.J., and D.B. designed the research. I.V. implemented the three-body fusion method in software, designed proteins, and performed biophysical characterization. Q.Y and Y.-J.P. performed cryo-EM and analysis of DARPin-grafted assemblies. Q.Y., S.J.W., and I.V. assembled and purified scaffold-target complexes. Y.-J.P., A.C., and I.V. performed negative-stain EM and analysis of base scaffolds. A.N., F.B., A.S. collected and analyzed mass spectrometry data. H.M. provided the DARPin sequence, information, and HSA protein. D.D.S., G.U., J.A.F., Y.H. assisted with experiments and scientific discussions. R.A.L. contributed the rop20 dimer scaffold. A.P., V.H.W., G.J.J., D.V., and D.B. supervised the research. All authors discussed results and commented on the manuscript.

## Acknowledgments

Research reported in this publication was supported by the National Institute Of General Medical Sciences, by the National Institutes of Health under Award Number T32GM008268 to I.V., as well as the Open Philanthropy Project, Howard Hughes Medical Institute, and NSF Grant CHE-1629214 to D.B. NIH grant under award AI150464 provided support to G.J.J. Cryo-EM work under G.J.J. was performed in the Caltech Beckman Institute Resource Center for Transmission Electron Microscopy. We also thank Dr. Songye Chen and Dr. Andrey Malyutin at Caltech for technical assistance. This work was also supported by the NIAID / NIH (HHSN272201700059C to D.V.) NIGMS / NIH (R01GM120553 to D.V.), a Pew Biomedical Scholars Award to D.V., and a Burroughs Wellcome Investigators in the Pathogenesis of Infectious Diseases award to D.V. This work was also supported by NIH grant P41GM128577 to V.H.W and Swiss National Science Foundation grant 310030_192689 to A.P. In addition, we thank Kathryn Burnett and Greg Hura for SAXS data collection through the SIBYLS mail-in SAXS program at the Advanced Light Source (ALS), a national user facility operated by Lawrence Berkeley National Laboratory on behalf of the Department of Energy, Office of Basic Energy Sciences, through the Integrated Diffraction Analysis Technologies (IDAT) program, supported by DOE Office of Biological and Environmental Research. Additional support comes from the National Institute of Health project ALS-ENABLE (P30 GM124169) and a High-End Instrumentation Grant S10OD018483. A.C. is a recipient of the Human Frontiers Science Program Long Term Fellowship. A.C. and D.D.S. received Washington Research Foundation fellowships. We thank Albumedix for providing high quality Veltis-grade HSA. I.V. thanks Shane Caldwell (University of Washington) for discussion on SAXS analysis and Vikram Mulligan (Flatiron Institute) for general assistance with RosettaScripts.

## References

1. Y Liu, DT Huynh, TO Yeates. A 3.8 Å resolution cryo-EM structure of a small protein bound to an imaging scaffold. Nat Commun. 10, 1864. (2019) doi: 10.1038/s41467-019-09836-0

2. Q Yao, SJ Weaver, JY Mock, GJ Jensen. Fusion of DARPin to Aldolase Enables Visualization of Small Protein by Cryo-EM. Structure. 27, 1148–1155.e3. (2019) doi: 10.1016/j.str.2019.04.003

3. J Marcandalli, B Fiala, S Ols, et al. Induction of Potent Neutralizing Antibody Responses by a Designed Protein Nanoparticle Vaccine for Respiratory Syncytial Virus. Cell. 176, 1420–1431.e17. (2019) doi: 10.1016/j.cell.2019.01.046

4. G Ueda, A Antanasijevic, J.A. Fallas et al. Tailored Design of Protein Nanoparticle Scaffolds for Multivalent Presentation of Viral Glycoprotein Antigens. bioRxiv (2019) doi: 10.1101/2020.01.29.923862

5. JE Padilla, C Colovos, TO Yeates. Nanohedra: using symmetry to design self assembling protein cages, layers, crystals, and filaments. Proc Natl Acad Sci U S A. 98, 2217–2221. (2001) doi: 10.1073/pnas.041614998

6. JC Sinclair, KM Davies, C Vénien-Bryan, ME Noble. Generation of protein lattices by fusing proteins with matching rotational symmetry. Nat Nanotechnol. 6, 558–562. (2011) doi: 10.1038/nnano.2011.122

7. YT Lai, E Reading, GL Hura, et al. Structure of a designed protein cage that self-assembles into a highly porous cube. Nat Chem. 6, 1065–1071. (2014) doi: 10.1038/nchem.2107

8. YT Lai, D Cascio, TO Yeates. Structure of a 16-nm cage designed by using protein oligomers. Science. 336, 1129. (2012) doi: 10.1126/science.1219351

9. S Badieyan, A Sciore, JD Eschweiler, et al. Symmetry-Directed Self-Assembly of a Tetrahedral Protein Cage Mediated by de Novo-Designed Coiled Coils. Chembiochem. 18, 1888–1892. (2017) doi: 10.1002/cbic.201700406

10. A Sciore, M Su, P Koldewey, et al. Flexible, symmetry-directed approach to assembling protein cages. Proc Natl Acad Sci U S A. 113, 8681–8686. (2016) doi: 10.1073/pnas.1606013113

11. AS Cristie-David, J Chen, DB Nowak, et al. Coiled-Coil-Mediated Assembly of an Icosahedral Protein Cage with Extremely High Thermal and Chemical Stability. J Am Chem Soc. 141, 9207–9216. (2019) doi: 10.1021/jacs.8b13604

12. KA Cannon, VN Nguyen, C Morgan, TO Yeates. Design and Characterization of an Icosahedral Protein Cage Formed by a Double-Fusion Protein Containing Three Distinct Symmetry Elements. ACS Synth Biol. 9, 517–524. (2020) doi: 10.1021/acssynbio.9b00392

13. NP King, W Sheffler, MR Sawaya, et al. Computational design of self-assembling protein nanomaterials with atomic level accuracy. Science. 336, 1171–1174. (2012) doi: 10.1126/science.1219364

14. NP King, JB Bale, W Sheffler, et al. Accurate design of co-assembling multi-component protein nanomaterials. Nature. 510, 103–108. (2014) doi: 10.1038/nature13404

15. S Gonen, F DiMaio, T Gonen, D Baker. Design of ordered two-dimensional arrays mediated by noncovalent protein-protein interfaces. Science. 348, 1365–1368. (2015) doi: 10.1126/science.aaa9897

16. JB Bale, S Gonen, Y Liu, et al. Accurate design of megadalton-scale two-component icosahedral protein complexes. Science. 353, 389–394. (2016) doi: 10.1126/science.aaf8818

17. Y Hsia, JB Bale, S Gonen, et al. Design of a hyperstable 60-subunit protein dodecahedron. Nature. 535, 136–139. (2016) doi: 10.1038/nature18010

18. JA Fallas, G Ueda, W Sheffler, et al. Computational design of self-assembling cyclic protein homo-oligomers. Nat Chem. 9, 353–360. (2017) doi: 10.1038/nchem.2673

19. YT Lai, KL Tsai, MR Sawaya, FJ Asturias, TO Yeates. Structure and flexibility of nanoscale protein cages designed by symmetric self-assembly. J Am Chem Soc. 135, 7738–7743. (2013) doi: 10.1021/ja402277f

20. PA Kratz, B Böttcher, M Nassal. Native display of complete foreign protein domains on the surface of hepatitis B virus capsids. Proc Natl Acad Sci U S A. 96, 1915–1920. (1999) doi: 10.1073/pnas.96.5.1915

21. F Coscia, LF Estrozi, F Hans, et al. Fusion to a homo-oligomeric scaffold allows cryo-EM analysis of a small protein. Sci Rep. 6, 30909. (2016) doi: 10.1038/srep30909

22. Y Liu, S Gonen, T Gonen, TO Yeates. Near-atomic cryo-EM imaging of a small protein displayed on a designed scaffolding system. Proc Natl Acad Sci U S A. 115, 3362–3367. (2018) doi: 10.1073/pnas.1718825115

23. A Plückthun. Designed ankyrin repeat proteins (DARPins): binding proteins for research, diagnostics, and therapy. Annu Rev Pharmacol Toxicol. 55, 489–511. (2015) doi: 10.1146/annurev-pharmtox-010611-134654

24. S Hansen, JC Stüber, P Ernst, et al. Design and applications of a clamp for Green Fluorescent Protein with picomolar affinity. Sci Rep. 7, 16292. (2017) doi: 10.1038/s41598-017-15711-z

25. A Batyuk, Y Wu, A Honegger, MM Heberling, et al. DARPin-Based Crystallization Chaperones Exploit Molecular Geometry as a Screening Dimension in Protein Crystallography. J Mol Biol. 428, 1574–1588. (2016) doi: 10.1016/j.jmb.2016.03.002

26. Y Wu, A Batyuk, A Honegger, et al. Rigidly connected multispecific artificial binders with adjustable geometries. Sci Rep. 7, 11217. (2017) doi: 10.1038/s41598-017-11472-x

27. RP Watson, MT Christen, C Ewald, et al. Spontaneous self-assembly of engineered armadillo repeat protein fragments into a folded structure. Structure. 22, 985–995. (2014) doi: 10.1016/j.str.2014.05.002

28. L Doyle, J Hallinan, J Bolduc, et al. Rational design of α-helical tandem repeat proteins with closed architectures. Nature. 528, 585–588. (2015) doi: 10.1038/nature16191

29. E Michel, A Plückthun, O Zerbe. Peptide-Guided Assembly of Repeat Protein Fragments. Angew Chem Int Ed Engl. 57, 4576–4579. (2018) doi: 10.1002/anie.201713377

30. CE Correnti, JP Hallinan, LA Doyle, et al. Engineering and functionalization of large circular tandem repeat protein nanoparticles. Nat Struct Mol Biol. 27, 342–350. (2020) doi: 10.1038/s41594-020-0397-5

31. TJ Brunette, F Parmeggiani, PS Huang, et al. Exploring the repeat protein universe through computational protein design. Nature. 528, 580–584. (2015) doi: 10.1038/nature16162

32. SE Boyken, Z Chen, B Groves, et al. De novo design of protein homo-oligomers with modular hydrogen-bond network-mediated specificity Science. 352, 680–687. (2016) doi: 10.1126/science.aad8865

33. DN Mastronarde. Automated electron microscope tomography using robust prediction of specimen movements. J Struct Biol. 152, 36–51. doi: 10.1016/j.jsb.2005.07.007

34. SQ Zheng, E Palovcak, J-P Armache, et al. bioRxiv 061960; (2005) doi: 10.1101/061960

35. T Grant, N Grigorieff. Measuring the optimal exposure for single particle cryo-EM using a 2.6 Å reconstruction of rotavirus VP6. Elife. 4, e06980. (2015) doi: 10.7554/eLife.06980

36. K Zhang. Gctf: Real-time CTF determination and correction. J Struct Biol. 193, 1–12. (2016) doi: 10.1016/j.jsb.2015.11.003

37. J Zivanov, T Nakane, BO Forsberg, et al. New tools for automated high-resolution cryo-EM structure determination in RELION-3. Elife. 7, e42166. (2018) doi: 10.7554/eLife.42166

38. A Punjani, JL Rubinstein, JL Fleet, et al. cryoSPARC: algorithms for rapid unsupervised cryo-EM structure determination. Nat Methods. 14, 290–296. (2017) doi: 10.1038/nmeth.4169

39. D Asarnow, E Palovcak, Y Cheng. UCSF pyem v0.5. Zenodo (2019) doi: 10.5281/zenodo.3576630

40. EF Pettersen, TD Goddard, CC Huang, et al. UCSF Chimera--a visualization system for exploratory research and analysis. J Comput Chem. 25, 1605–1612. (2004) doi: 10.1002/jcc.20084

41. P Emsley, K Cowtan. Coot: model-building tools for molecular graphics. Acta Crystallogr D Biol Crystallogr. 60, 2126–2132. (2004) doi: 10.1107/S0907444904019158

42. A Kucukelbir, FJ Sigworth, HD Tagare. Quantifying the local resolution of cryo-EM density maps. Nat Methods. 11, 63–65. (2014) doi: 10.1038/nmeth.2727

43. D Tegunov, P Cramer. Real-time cryo-electron microscopy data preprocessing with Warp. Nat Methods. 16, 1146–1152. (2019) doi: 10.1038/s41592-019-0580-y

44. ZL VanAernum, F Busch, BJ Jones, et al. Rapid online buffer exchange for screening of proteins, protein complexes and cell lysates by native mass spectrometry. Nat Protoc. 15, 1132–1157. (2020) doi: 10.1038/s41596-019-0281-0

45. M van de Waterbeemd, KL Fort, D Boll, et al. High-fidelity mass analysis unveils heterogeneity in intact ribosomal particles. Nat Methods. 14, 283–286. (2017) doi: 10.1038/nmeth.4147

46. KL Fort, M van de Waterbeemd, D Boll, et al. Expanding the structural analysis capabilities on an Orbitrap-based mass spectrometer for large macromolecular complexes. Analyst. 143, 100–105. (2017) doi: 10.1039/c7an01629h

47. ZL VanAernum, JD Gilbert, ME Belov, et al. Surface-Induced Dissociation of Noncovalent Protein Complexes in an Extended Mass Range Orbitrap Mass Spectrometer. Anal Chem. 91, 3611–3618. (2019) doi: 10.1021/acs.analchem.8b05605

48. MT Marty, AJ Baldwin, EG Marklund, et al. Bayesian deconvolution of mass and ion mobility spectra: from binary interactions to polydisperse ensembles. Anal Chem. 87, 4370–4376. (2015) doi: 10.1021/acs.analchem.5b00140

49. FW Studier. Protein production by auto-induction in high density shaking cultures. Protein Expr Purif. 41, 207–234. (2005) doi: 10.1016/j.pep.2005.01.016

50. T Grant, A Rohou, N Grigorieff. *cis*TEM, user-friendly software for single-particle image processing. Elife. 7, e35383. (2018) doi: 10.7554/eLife.35383

51. D Schneidman-Duhovny, M Hammel, JA Tainer, Sali A. Accurate SAXS profile computation and its assessment by contrast variation experiments. Biophys J. 105, 962–974. (2013) doi: 10.1016/j.bpj.2013.07.020

52. D Schneidman-Duhovny, M Hammel, JA Tainer, A Sali. FoXS, FoXSDock and MultiFoXS: Single-state and multi-state structural modeling of proteins and their complexes based on SAXS profiles. Nucleic Acids Res. 44, W424–W429. (2016) doi: 10.1093/nar/gkw389

