## Supplementary material for "Generation of ordered protein assemblies using rigid three-body fusion": fusion_supplement

Supplementary Figures

- Figure S1 - Size exclusion chromatography
- Figure S2 - Marginal designs
- Figure S3 - D2-1.4H + GFP-DARPin alignment based grafting
- Figure S4 - Native mass spectrometry, round-1 designs
- Figure S5 - Native mass spectrometry, round-1 designs
- Figure S6 - ankyrin/DARPin homo-dimer orientation selection criteria
- Figure S7 - Cryo-EM of D2-1.4H-GFP.v1 core and D2-21.8.GFP.v2

Supplementary Tables

- Supplementary Table 1 - Original design sequences
- Supplementary Table 2 - Building block table for successful designs
- Supplementary Table 3 - GFP and HSA-binding variant sequences
- Supplementary Table 4 - Building blocks with crystal structures
- Supplementary Table 5 - Native-MS expected and determined masses

Supplementary Text Files

- Supplementary Text File 1 - RosettaScript design protocol
- Supplementary Text File 2 - D2 symmetry definition
- Supplementary Text File 2 - D3 symmetry definition

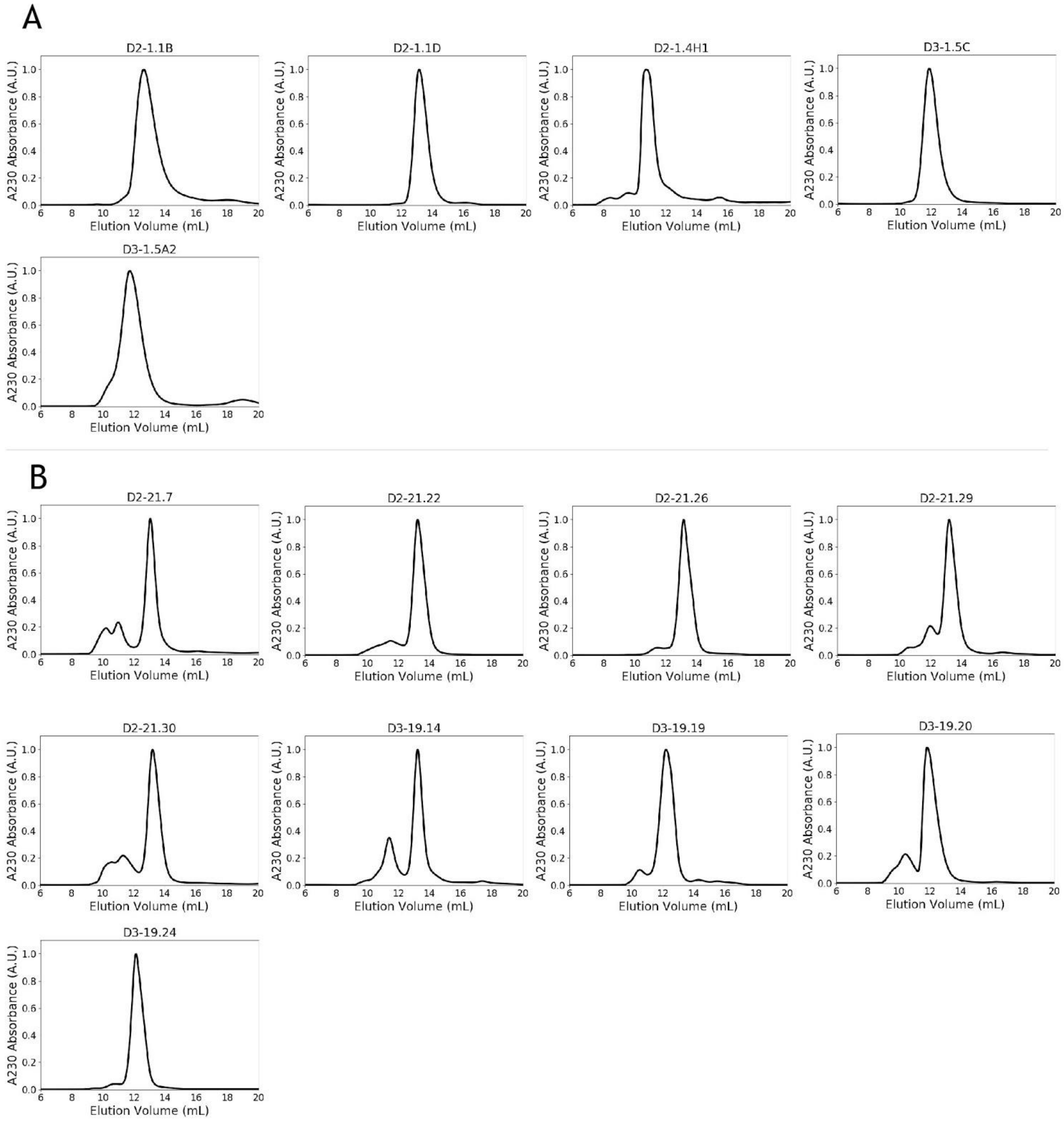


**Supplemental Figure 1**: Size exclusion chromatography of post-IMAC eluate on a Superdex 200 Increase 10/300 GL (not re-chromatography) shows a dominant species for constructs from the (A) first and (B) second design rounds.

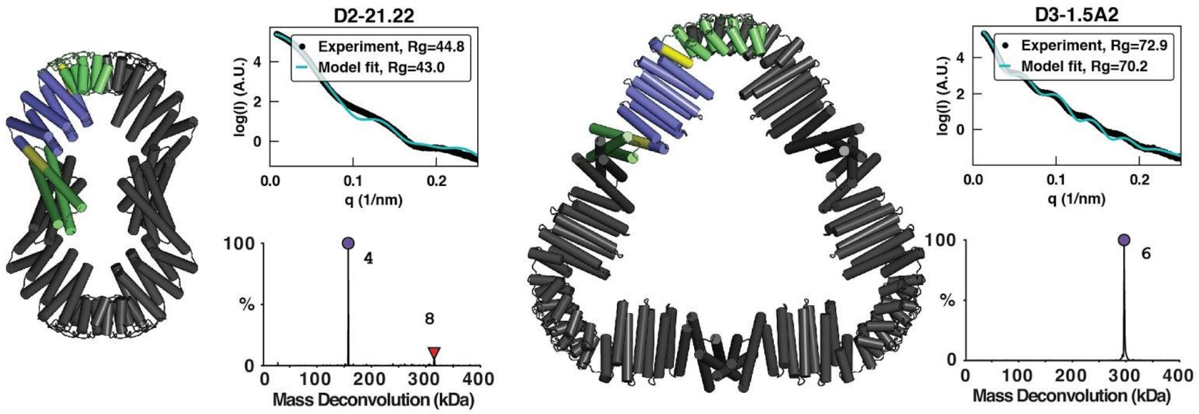


**Supplemental Figure 2:** Two designs showed promising results according to SAXS and native-MS, but appeared disordered by EM. It may be that the assemblies are sensitive to the low pH of uranyl formate stain or are simply more unstable. Consistent with lesser stability for design D3-1.5A2, incomplete assembly was observed when using offline buffer exchange for this design, indicating complex dissociation and/or unfolding as a result of extended protein storage in sub-optimal buffer (AmAC) that occurred between offline buffer exchange and native-MS measurement. Along these lines, no complex dissociation was observed when using online buffer-exchange MS (which was used to generate all the data shown) as the time between buffer-exchange to AmAc and native-MS measurement is drastically reduced with this method. As online buffer exchange was a newer and improved protocol, it was never used with D2-21.22 or other round-2 designs.

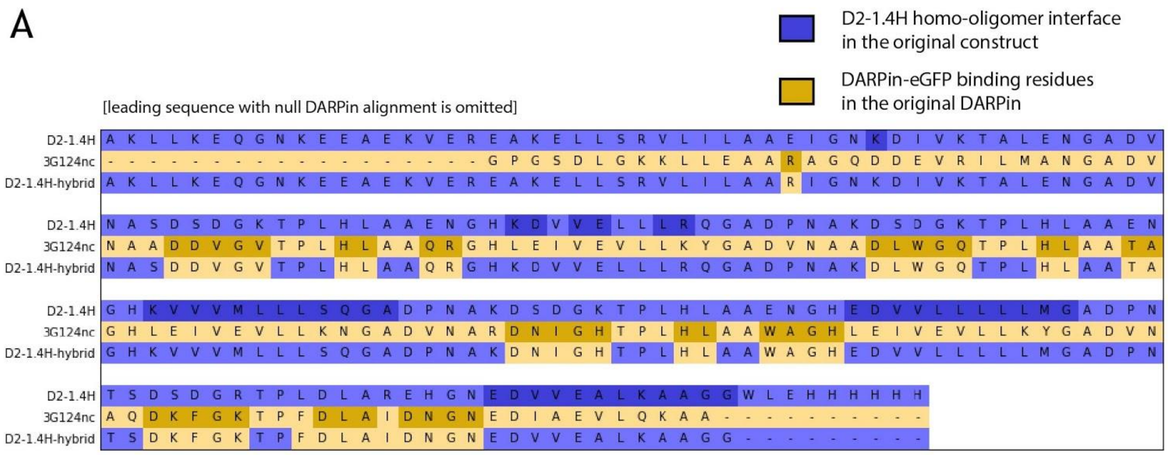


**Supplemental Figure 3:** Alignment-based construction of a hybrid DARPin scaffold “D2-1.4H.GFP.v1”**.** An alignment is performed during DARPin grafting to ensure that residues responsible for homo-oligomerization in the base construct (D2-1.4H) are preserved after hybridization with the DARPin. Binding residues (D2-1.4H homo-oligomerization and the DARPin-GFP interface) in the source constructs are darkened and the hybrid construct is colored according to whichever source construct the sequence was based on.

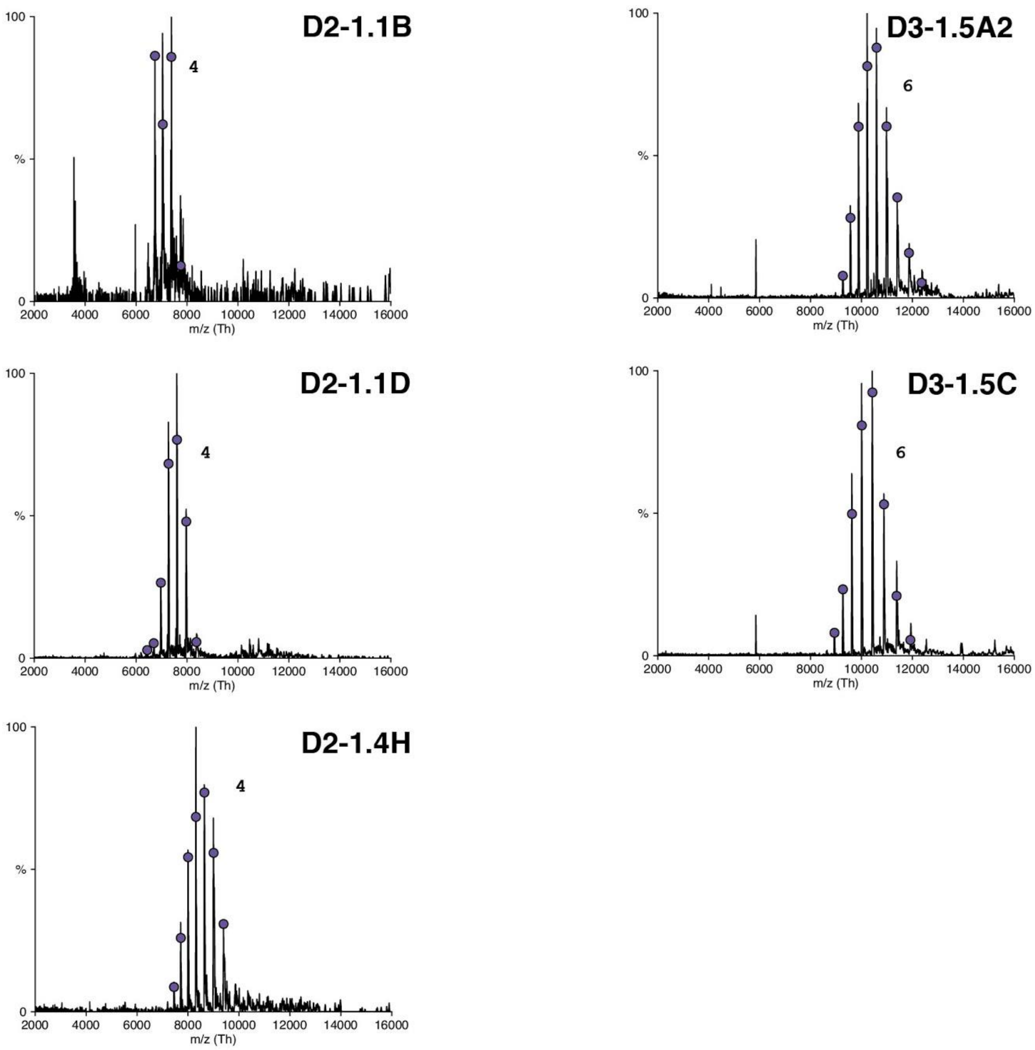


**Supplementary Figure 4**: Raw spectra showing relative abundance vs *m/z* of the first-round designs. The charge state distributions are labeled with purple circles and the oligomeric states are noted.

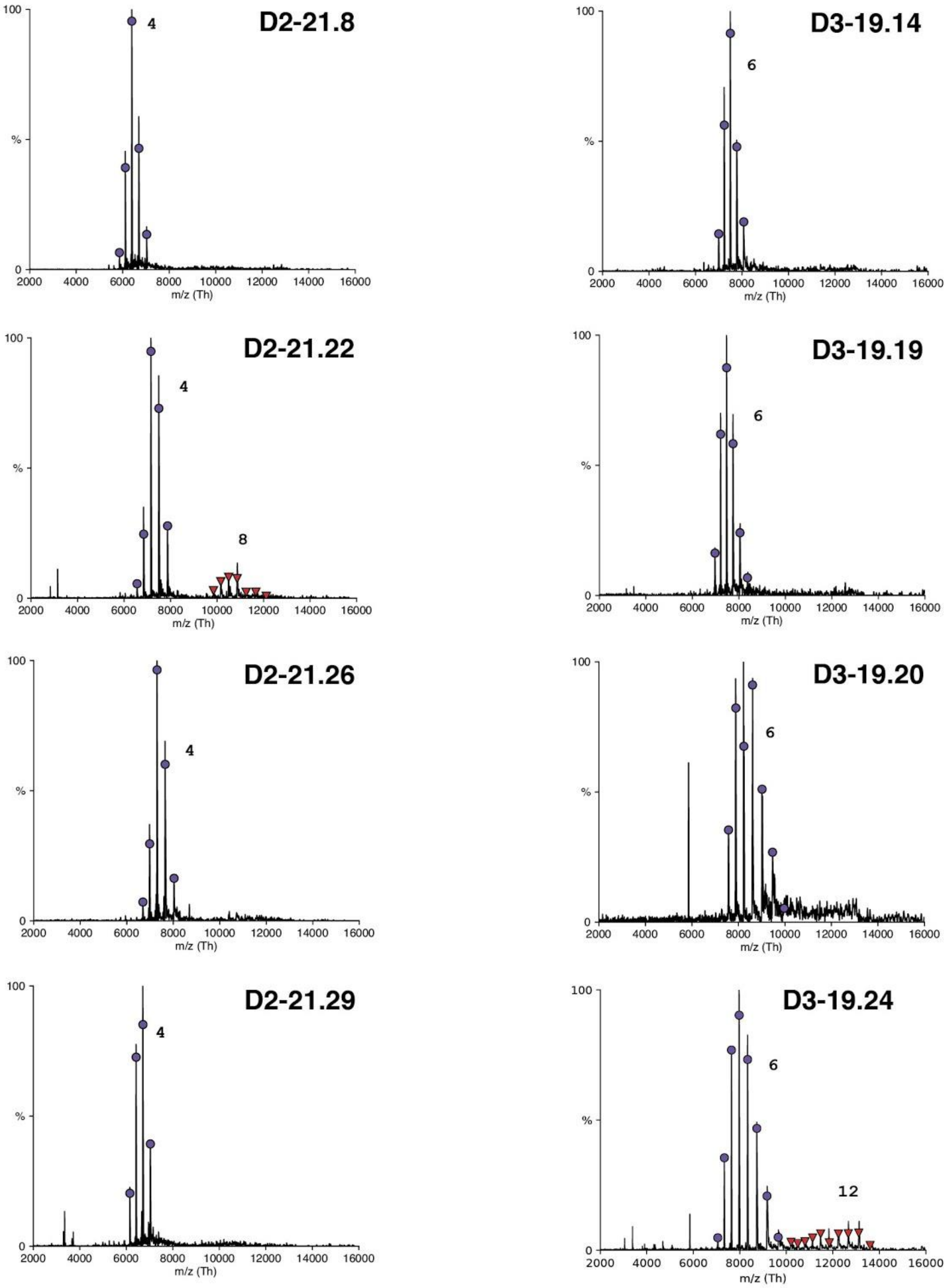


**Supplementary Figure 5**: Raw spectra showing relative abundance vs m/z of the second-round designs. The charge state distributions are labeled with purple circles and the oligomeric states are noted. Designed oligomers D2-21.22 and D3-19.24 appear to show low levels of self-association.

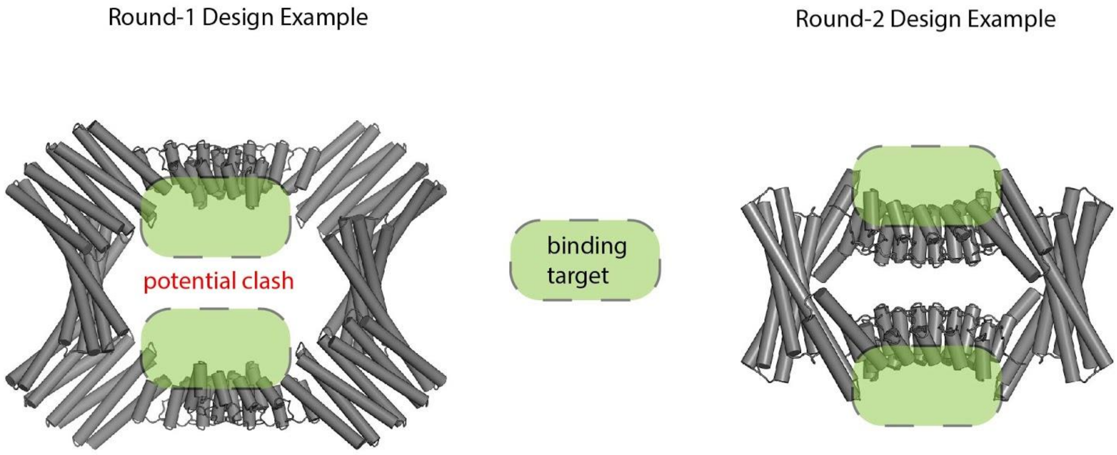


**Supplementary Figure 6**. Orientation criteria for the ankyrin/DARPin homo-dimer that were applied in the second design round as illustrated. Target-binding by DARPins usually occurs at the concave surface between loops and helices and it was thought that the flipped ankyrin/DARPin homo-dimer orientation in the round-2 designs (right) would generally orient binding-target copies away from one another.

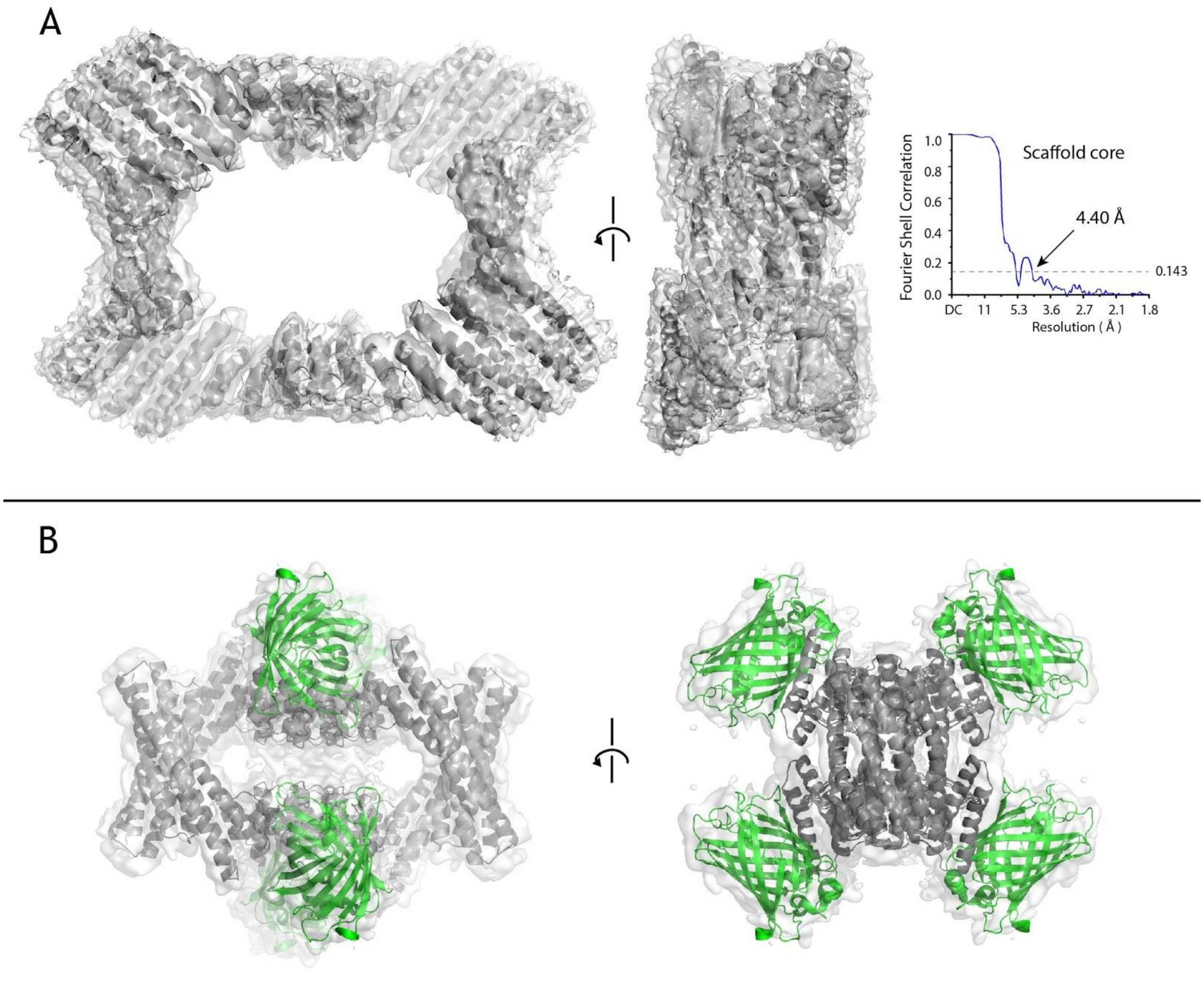


**Supplementary Figure 7**. Cryo-EM for GFP-related scaffolds with embedded models. A) The scaffold core, resolved at 4.38 Å based on the FSC=0.143 criterion, determined from the full assembly of GFP with scaffold D2-1.4H.GFP.v1, with GFP masked. B) The full assembly of GFP with second-round design D2-21.8.GFP.v2, resolved at approximately 6 Ångströms.

**Supplementary Table 1**: List of sequences whose SEC, nMS, and SAXS data were in agreement with the corresponding design model. Underlined linker and (His)6-tag denote a region added after computational design and not modeled during SAXS analysis.

| Round-1 Design Sequences |
| --- |
| D2-1.1B<br><br>MASDYLRLATEHNKLATEAASLAAELAASAVTLTVTDPSKTAQEHTELASRLLEMMSQFLKA<br>AQELTREAIRKEGRNEESEKTLRKSKSSYKALKALLKAIKAIEKGDVETAVRAAQEAVRLASE<br>AGNNNVLRAVAEVALAIAKVAEEQGNVEVAVKAAQVAVSAALNAGDEDVLKKVAEQASRISK<br>EAEKQGNQEVSKKALSVSLIAAAASGDKDLVKDLLESGADVNASSSDGKTPLHVAAENGHA<br>KVVLLLLEQGADPNAKDSDGKTPLHLAAENGHAVVVALLLMHGADPNAKDSDGKTPLHLAA<br>ENGHEEVVILLLAMGADPNTSDSDGRTPLDLAREHGNEEVVKVLEDHGGWLEHHHHHH |
| D2-1.1D<br><br>MASEKARIAVENLEAALRLNKAAIEMAKSAIKITRDNSSDEKATRYSLLTAKVLVMSLELLTASL<br>ELAEKALREEGSDDSAEKVRKEAEEILSKAVEEAVRVMQEMVTIMKRTGSNDSLREVAELAL<br>RVAKAAEKAGNVEVAVQAARVAVEAAKQAGDNDVLRKVAEQALRIAKEAEKQGNVEVAVKA<br>AKVAVEAAKQAGDEDVLKKVAEQASRIASEASKQGNKEVASKALIVAAQAGSKEAVKKAIES<br>GADVNASDSDGRTPLHHAAENGHAEVVALLIEKGADVNAKDSDGRTPLHHAAENGHDEVVL<br>ILLLKGADVNAKDSDGRTPLHHAAENGHKRVVLVLILAGADVNTSDSDGRTPLDLAREHGNE<br>EVVKALEKQGGWLEHHHHHH |
| D2-1.4H<br><br>MGSEKARIAVENLEAALRLNRAAAEMQKSAIKIMDDNSDDEKALRYLRLTTKVLRMSVELLRA<br>SLELAEKALREEGSDDSAEKVRKEAEEILKESTAILKLADAATKVADIKHDIKKAKEQQEQGNK<br>EEAEKTLREATEKIKRVTEELEKIAKNSKTPEIALKAAEALVKLIKLLIEIAKLLQEQGNKEEAEK<br>VLREATELIKRVTELLEKIAKNSDTPELALRAAELLVRLIKLLIEIAKLLQEQGNKEEAEKVLREA<br>TELIKRVTELLEKIAKNSDTPELASRAAELLVRLIKLLQEIAKLLKEQGNKEEAEKVEREAKELL<br>SRVLILAAEIGNKDIVKTALENGADVNASDSDGKTPLHLAAENGHKDVVELLLRQGADPNAKD<br>SDGKTPLHLAAENGHKVVVMLLLSQGADPNAKDSDGKTPLHLAAENGHEDVVLLLLLMGAD<br>PNTSDSDGRTPLDLAREHGNEDVVEALKAAGGWLEHHHHHH |
| D3-1.5C<br><br>MGSEKARIAVENLEAALRLNRAAAEMQKSAAKIVADNASDEKALRYLRLTTKVLRMSVELLRA<br>SLELAEKALREEGSDDSAEKVRKEAEEILKESTEILKEADQITEVADLAFELANKATDEELRKEI<br>SKCARLALELASRSTNDELIKQILEVAKLAFELASKATDEELIKLILKCCQAAFERASRSTNDEE<br>IKKILEVAKRAFETASKATDEEEIKSILLICAAALGNKDAVKSAIENGADVNASDSDGRTPLHHA<br>AENGNAEVVALLIEKGADVNAKDSDGRTPLHHAAENGHDEVVLILLLKGADVNAKDSDGRTP<br>LHHAAENGHKRVVLVLILAGADVNTSDSDGRTPLDLAREHGNEEVVKALEKQGGWLEHHHH<br>HH |

D3-1.5A2

MGSEKARIAVENLEAALRLNRAAAEMQKSAAKIVADNASDEKALRYLRLTTKVLRMSVELLRA
SLELAEKALREEGSDDSAEKVRKEAEEILKESTEILKLADQITEVADLAFELANKATDEELRKEI
VKCAKLALELASRSTNDELKKQILEVAKLAFELASKATDEELIKEILKCCQLAFELASRSTNDEL
IKQILEVAKLAFELASKATDEELIKEILKCCQLAFELASRSTNDELIKLILEVAKAAFERASKATD
EEEIKEILKVCQEAFEEASRSTNDEEIASILLVAAALLGNKDAVKDAIENGADVNASDSDGRTP
LHHAAENGNAEVVALLIEKGADVNAKDSDGRTPLHHAAENGHDEVVLILLLKGADVNAKDSD
GRTPLHHAAENGHKRVVLVLILAGADVNTSDSDGRTPLDLAREHGNEEVVKALEKQGGWLE
HHHHHH

Round-2 Design Sequences (D2 symmetry)

D2-21.6

MHHHHHHGSGSALEKIAKLIIEAARLSAELARRAARASAEMARLAIEAVSKERGSSELLKIVAD
LIVEAQEAVVRLIIESQQIAAKLAEDLIRAAKEAASDESKMEEVAKEVQERAERAARDIEEKLLK
VLIELIKKLARSIGDEERLKATKLAEEAIRVAREVGDSLLERIALEAALKGDSRAAKAVLKAAEL
AREAAERGDEEKVKAAALIAAAAAGDKDAVKDLIENGADVNGRDSDGRTPLHHAAENGNAE
VVALLIAKGADVNAKDSDGRTPLHHAAENGHEEVVLILLLKGADVNAKDSDGRTPLHHAAEN
GHKRVVLVLILAGADVNTSDSDGRTPLDLAREHGNEEVVKALEKQ

D2-21.8

MHHHHHHGSGSEKARIAVENLEAALRLNRAAAEMQKSAIKIMRDNSSDEKAFRYLLLTTKVL
KMSVELLRASLELAEKALREEGSDDSAEKVRKEAEEILKESTEILKRAELETLKAAVRVAAEAA
ARNATDEEERKRIEEELKKAEERANRSTNEEEIKKILEEALARFLIILAEKGAKEAVKLALEAGA
DVNGKDSDGKTPLHLAAENGHAKVVLLLLEQGADPNAKDSDGKTPLHLAAENGHAVVVALL
LMHGADPNAKDSDGKTPLHLAAENGHEEVVILLLAMGADPNTSDSDGRTPLDLAREHGNEE
VVKVLEDHG

D2-21.22

MHHHHHHGSGSEKARIAVENLEAALRLNRAAAEMQLSAAKIAADNFSDKKAAEYTRLTTKVL
EMSVELLRASLELAEKALREEGSDDSAEKVRKEAEEILKESKEILEAAEALTRIAHLARKAAES
TDPEEALKIAKEAIEIALKTVKENPSELALQAVLAAVILASAVAKRVTDPDKALKIAKLVIELALEA
VKEDPSTDALRAVLEAVRLAEEVARRVTDPIKALKIAALVIQLAAEAAKEDPSEEAQRALKLAA
ELAAEALERGADVNYHDEDGRTPLHHAAEAGADEAVLILLLKGADVNAKDSDGRTPLHHAAE
NGNKRVVLVLILAGADVNTSDSDGRTPLDHAREHGNEEVVKALEKQ

D2-21.26

MHHHHHHGSGSEKARIAVENLEAALRLNRAAAEMQASAAKIVADNTSDEKAYRYLELTTKVL
LMSVELLRASLELAEKALREEGSDDSAEKVRKEAEEILKESTLALEAAEELTKAAKAALRARE

ASERGDEEEFRKAAEEALEAAKRVVERAKKAGIPELVAAAAAVALAIAELAAKNGDKEVFKKA
AESALEVAKRLVEVASKEGDPELVLEAAKVALRVAELARKNGDKEVLKKAALSAAEVALRLAE
VAKKEGDPDLVREAAEILADALEKGLDVNIHDEDGRTPLHHAAELGADEAVLILLLAGADVNA
KDSDGRTPLHHAAENGHKRVVLVLILAGADVNTSDSDGRTPLDHAREHGNEEVVKALEKQ

D2-21.29

MHHHHHHGSGSEKARIAVENLEAALRLNRAAAEMQKSAIKIADDNRSDEKALRYALLTTKVL
EMSVELLRASLELAEKALREEGSDDSAEKVRKEAEEILEKSSRILAEAFVITARLATELARLLQ
EKAKKTGDAKELREAKRALKEAAEYVEKALKINKDDDEARELLERIEEELKKVEKLLEEILIKAA
ARGDKDLVKLALKAGADVNASDSDGKTPLHKAAENGHAKVVLLLLEQGADPNAKDSDGKTP
LHLAAENGHAVVVALLLMHGADPNAKDSDGKTPLHLAAENGHEEVVILLLAMGADPNTSDSD
GRTPLDLAREHGNEEVVKVLEDHG

D2-21.30

MHHHHHHGSGSEKARIAVENLEAALRLNRAAAEMQKSAIKIAEDNSSDEKAIRYTLLTTRVLE
MSFELLRASLELAEKALREEGSDDSAEKVRKEAEEILEESLRILAEAFVRTARFLKELAERLQE
RAKKTGDPELLAEAYEALREAVEFVKKAEKINPDNERAKKTLEELKEELRKVEELLKELLIRAA
ERGDKDTVRRALEAGADVNAKDSDGKTPLHLAAENGHAKVVLLLLEQGADPNAKDSDGKTP
LHLAAENGHAVVVALLLMHGADPNAKDSDGKTPLHLAAENGHEEVVILLLAMGADPNTSDSD
GRTPLDLAREHGNEEVVKVLEDHG

Round-2 Design Sequences (D3 symmetry)

D3-19.14

MHHHHHHGSSEKARIAVENLEAALRLNRAAAEMQKSAIKIARDNRSDKKALLYLLLATYVLEM
SLELLRASLELAEKALREEGSDDSAEKVRKEAEEILKESKEIFLRAALETAKAAAEYVEEAARE
AERRGNPELRDAAKALRKYLEEANEEAAKQGNAEKILRVALAALLIAAAALGDKDLVKDLIEM
GADVNGHDLDGRTPLHLAAAAGHAEVVALLIEKGADVNAKDSDGRTPLHHAAENGHDEVVLI
LLLKGADVNAKDSDGRTPLHHAAENGHKRVVLVLILAGADVNTSDSDGRTPLDLARENGNEE
VVKVLEKA

D3-19.19

MHHHHHHGSSEKARIAVENLEAALRLNRAAAEMQKSAIKIVDDNSSDVRAIEYLALTSAVLAE
SLLLLLASLELAEKALREEGSDDSAEKVRKEAEEILEESARIAAEAAEESLRAAEEAIELARKTG
DSDALRAAAEALKAARAAVRAAIAANPDDDKAEEIAKRLEEALNRVLHEAAERGDQDAVKLVI
EAGGDVNARDSDGRTPLHHAAENGHAEVVALLIRKGADVNAKDSDGRTPLHHAAENGHDE
VVLILLLKGADVNAKDSDGRTPLHHAAENGHKRVVLVLILAGADVNTSDSDGRTPLDHAREN
GNEKVVKALQEQ

| |
|---|
| D3-19.20<br><br>MHHHHHHGSSEKARIAVENLEAALRLNRAAAEMQKSAIKIALDNSSDEKAIRYARLTTKVLKM<br>SVELLRASLELAEKALREEGSDDSAEKVRKEAEEILKESTLILEAADLATALLDLLQKVRKVEK<br>EIKSNKDNEEAVETAARLAIELARVAKRLEELAKKLGDGFLKKLAEKAIKIAARALEVALEAGYD<br>VNAKDSDGRTVLHHAAENGALEVVLLALLNGADVNAKDSDGRTPLHHAAENGNKRVVLVLIL<br>AGADVNTSDSDGRTPLDLARENGNEEVVKALERR |
| D3-19.24<br><br>MHHHHHHGSSEKARIAVENLEAALRLNRAAAEMQFLAIKIMLLNSSDEKAARFLRLTTKVLKM<br>SVELLRASLELAEKALREEGSDDSAEKVRKEAEEILKESTEILEATEEATKLLELLEEARKVEE<br>AIKSNPDNDEAVETAKRIAEEARKVALKLFEYASKLGIPLLAKAAAEALAVALKAGADPNAKDS<br>DGKTPLHHAAEAGVKLAVMLLLSHGADPNAKDSDGKTPLHLAAENGHEDVVLLLLLMGADP<br>NTSDSDGRTPLDLAREHGNEDVVKALKAAG |

**Supplementary Table 2**: Constituent building blocks for successful and marginal designs are shown. Those building blocks that have been SAXS-verified, but not crystallized, are colored green; most successful designs are composed with two of three components having only SAXS verification. Two successful designs were created without any building block crystal verification whatsoever.

| Design ID | N-terminal oligomer | Spacer | C-terminal oligomer |
|---|---|---|---|
| D2-1.1B | rop4 | DHR62 | ank3C21 |
| D2-1.1D | rop20 | DHR62 | ank1C2G3 |
| D2-1.4H | rop20 | DHR68 | ank3C22 |
| D3-1.5C | rop20 | DHR15 | ank1C2G3 |
| D3-1.5A2 (marginal) | rop20 | DHR15 | ank1C2G3 |
| D2-21.8 | rop20 | DHR15 | ank3C21 |
| D2-21.22 (marginal) | rop20 | DHR57 | ank1C2G3 |
| D2-21.29 and D2-21.30 | rop20 | DHR82 | ank3C21 |
| D2-21.26 | rop20 | DHR71 | ank1C2G3 |
| D3-19.19 | rop20 | DHR82 | ank1C2G3 |
| D3-19.14 | rop20 | DHR76 | ank1C2G3 |
| D3-19.20 | rop20 | DHR82 | ank1C2G3 |
| D3-19.24 | rop20 | DHR82 | ank3C22 |

**Supplementary Table 3**: GFP- and HSA-binding scaffold variants and the original HSA-DARPin sequence are listed, along with the corresponding resolution achieved or the observed failure mode, where applicable. Design IDs are comprised of the underlying scaffold ID plus the suffix v[#], to separate variants of the same scaffold, which were produced by shifting the grafted residues up or down by one or more ankyrin repeats.

D2-1.1D.GFP.v1: Aggregated

SEKARIAVENLEAALRLNKAAIEMAKSAIKITRDNSSDEKATRYSLLTAKVLVMSLELLTASLEL
AEKALREEGSDDSAEKVRKEAEEILSKAVEEAVRVMQEMVTIMKRTGSNDSLREVAELALRV
AKAAEKAGNVEVAVQAARVAVEAAKQAGDNDVLRKVAEQALRIAKEAEKQGNVEVAVKAAK
VAVEAAKQAGDEDVLKKVAEQASRIASEASKQGNKEVASKALIVAAQAGSKEAVKKAIESGA
DVNASDSDGRTPLHHAAENGHAEVVALLIEKGADVNAKDSNGHTPLLHAARNGHDEVVLILL
LKGADVNAKDDVGVTPLHLAAQRGHKRVVLVLILAGADVNTADLWGQTPLHLAATAGHLEVV
KALLKQGADVNARDNIGHTPLHLAAWAGHLEIVEVLLKYGADVNAQDKFGKTPFDLAIDNGN
EDIAEVLQKA

D2-1.4H.GFP.v1: 4.3 Å overall resolution

SEKARIAVENLEAALRLNRAAAEMQKSAIKIMDDNSDDEKALRYLRLTTKVLRMSVELLRASL
ELAEKALREEGSDDSAEKVRKEAEEILKESTAILKLADAATKVADIKHDIKKAKEQQEQGNKEE
AEKTLREATEKIKRVTEELEKIAKNSKTPEIALKAAEALVKLIKLLIEIAKLLQEQGNKEEAEKVL
REATELIKRVTELLEKIAKNSDTPELALRAAELLVRLIKLLIEIAKLLQEQGNKEEAEKVLREATE
LIKRVTELLEKIAKNSDTPELASRAAELLVRLIKLLQEIAKLLKEQGNKEEAEKVEREAKELLSR
VLILAARIGNKDIVKTALENGADVNASDDVGVTPLHLAAQRGHKDVVELLLRQGADPNAKDL
WGQTPLHLAATAGHKVVVMLLLSQGADPNAKDNIGHTPLHLAAWAGHEDVVLLLLLMGADP
NTSDKFGKTPFDLAIDNGNEDVVEALKAAGG

D2-1.5C.GFP.v1: Disordered

SEKARIAVENLEAALRLNRAAAEMQKSAAKIVADNASDEKALRYLRLTTKVLRMSVELLRASL
ELAEKALREEGSDDSAEKVRKEAEEILKESTEILKEADQITEVADLAFELANKATDEELRKEISK
CARLALELASRSTNDELIKQILEVAKLAFELASKATDEELIKLILKCCQAAFERASRSTNDEEIK
KILEVAKRAFETASKATDEEEIKSILLICAAALGNKDAVKSAIENGADVNASDSDGRTPLHHAA
ENGNAEVVALLIEKGADVNAKDSDGHTPLLHAARNGHDEVVLILLLKGADVNAKDDVGVTPL
HLAAQRGHKRVVLVLILAGADVNTADLWGQTPLHLAATAGHEEVVKALIKQGADVNARDNIG
HTPLHLAAWAGHLEIVEVLLKYGADVNAQDKFGKTPFDLAIDNGNEDIAEVLQKA

D2-21.8.GFP.v1: Slight aggregation

MHHHHHHGSGSEKARIAVENLEAALRLNRAAAEMQKSAIKIMRDNSSDEKAFRYLLLTTKVL
KMSVELLRASLELAEKALREEGSDDSAEKVRKEAEEILKESTEILKRAELETLKAAVRVAAEAA
ARNATDEEERKRIEEELKKAEERANRSTNEEEQKKILEEALGRFLIILARKGAKEAVKLALEAG
ADVNAADDVGVTPLHLAAQRGHAKVVLLLLEYGADPNAADLWGQTPLHLAATAGHAVVVAL
LLMHGADPNARDNIGHTPLHLAAWAGHEEVVILLLAMGADPNAQDKFGKTPLDLARDNGNE
EVVKVLEDHAA

| |
|---|
| D2-21.8.GFP.v2: 6-7 Å overall resolution with target and minor preferred orientation<br><br>MHHHHHHGSGSEKARIAVENLEAALRLNRAAAEMQKSAIKIMRDNSSDEKAFRYLLLTTKVL<br>KMSVELLRASLELAEKALREEGSDDSAEKVRKEAEEILKESTEILKRAELETLKAAVRVAAEAA<br>ARNATDEEERKRIEEELKKAEERANRSTNEEEQKKILEEALGRFLIILARKGAKEAVKLALEAG<br>ADVNAADDVGVTPLHLAAQRGHAKIVLLLLEYGADPNAADLWGQTPLHLAATAGHAVIVALLL<br>MHGADPNARDNIGHTPLHLAAWAGHEEIVILLLAMGADPNAQDKFGKTPLDLARDNGNEEV<br>VKVLEDHAA |
| D2-21.8.GFP.v3: Preferred orientation<br><br>MHHHHHHGSGSEKARIAVENLEAALRLNRAAAEMQKSAIKIMRDNSSDEKAFRYLLLTTKVL<br>KMSVELLRASLELAEKALREEGSDDSAEKVRKEAEEILKESTEILKRAELETLKAAVRVAAEAA<br>ARNATDEEERKRIEEELKKAEERANRSTNEEEQKKILEEALGRFLIILARKGAKEAVKLALEAG<br>ADVNAADDVGVTPLHLAAQRGHAKVVLLLLEQGADPNAADLWGQTPLHLAATAGHAVVVAL<br>LLMHGADPNARDNIGHTPLHLAAWAGHEEVVILLLAMGADPNAQDKFGKTPLDLARDNGNE<br>EVVKVLEDHAA |
| D2-21.29.GFP.v1: Severe aggregation<br><br>MHHHHHHGSGSEKARIAVENLEAALRLNRAAAEMQKSAIKIADDNRSDEKALRYALLTTKVL<br>EMSVELLRASLELAEKALREEGSDDSAEKVRKEAEEILEKSSRILAEAFVITARLATELARLLQ<br>EKAKKTGDAKELREAKRALKEAAEYVEKALKINKDDDEARELLERIEEELKKVEDLLGKILLEA<br>ARAGDKDLVKLALKAGADVNAADDVGVTPLHLAAQRGHAKVVLLLLEYGADPNAADLWGQT<br>PLHLAATAGHAVVVALLLMHGADPNARDNIGHTPLHLAAWAGHEEVVILLLAMGADPNAQDK<br>FGKTPLDLARDNGNEEVVKVLEDHAA |
| D2-21.29.GFP.v2: Preferred orientation and aggregated<br><br>MHHHHHHGSGSEKARIAVENLEAALRLNRAAAEMQKSAIKIADDNRSDEKALRYALLTTKVL<br>EMSVELLRASLELAEKALREEGSDDSAEKVRKEAEEILEKSSRILAEAFVITARLATELARLLQ<br>EKAKKTGDAKELREAKRALKEAAEYVEKALKINKDDDEARELLERIEEELKKVEKLLGEILLEA<br>ARAGDKDLVKLALKAGADVNAADDVGVTPLHLAAQRGHAKIVLLLLEYGADPNAADLWGQT<br>PLHLAATAGHAVIVALLLMHGADPNARDNIGHTPLHLAAWAGHEEIVILLLAMGADPNAQDKF<br>GKTPLDLARDNGNEEIVKVLEDHAA |
| D2-19.20.GFP.v1: Aggregated<br><br>MHHHHHHGSGSEKARIAVENLEAALRLNRAAAEMQKSAIKIALDNSSDEKAIRYARLTTKVLK<br>MSVELLRASLELAEKALREEGSDDSAEKVRKEAEEILKESTLILEAADLATALLDLLQKVRKVE<br>KEIKSNKDNEEAVETAARLAIELARVAKRLEELAKKLGDGFLKKLAEKAIKIAARALEVALEAGY<br>DVNAKDSDGATVLHHAARNGALEVVLLALLNGADVNAADDVGVTPLHLAAQRGNKRVVLVLI<br>LAGADVNAADLWGQTPLHLAATAGHLEVVKALLKRGADVNARDNIGHTPLHLAAWAGHLEIV<br>EVLLKYGADVNAQDKFGKTPFDLAIDNGNEDIAEVLQKA |

| |
|---|
| D2-21.8.HSA-C9.v2: 5.5 Å overall resolution with target<br><br>MHHHHHHSEKARIAVENLEAALRLNRAAAEMQKSAIKIMRDNSSDEKAFRYLLLTTKVLKMS<br>VELLRASLELAEKALREEGSDDSAEKVRKEAEEILKESTEILKRAELETLKAAVRVAAEAAARN<br>ATDEEERKRIEEELKKAEERANRSTNEEEIKKILEEALARFLLEAAWKGAKEAVKLALEAGAD<br>VNAADYFGHTPLHLAARNGHAKVVLLLLEQGADPNADDFAGSTPLHLAARAGHAVVVALLLM<br>HGADPNAVDSNGFTPLHLAAQKGHEEVVILLLAMGADPNAQDKFGKTPFDLAIDNGNEEVVK<br>VLEDHG |
| Original DARPin C9 (anti-HSA) - FLAG tag underlined<br><br>MRGSHHHHHHGSDLGKKLLEAAWWGQDDEVRILMANGADVNAADYFGHTPLHLAARNGH<br>LEIVEVLLKTGADVNADDFAGSTPLHLAARAGHLEIVEVLLKAGADVNAVDSNGFTPLHLAAQ<br>KGHLEIVEVLLKHGADVNAQDKFGKTPFDLAIDNGNEDIAEVLQKAAKLNDYKDDDDK |

**Supplementary Table 4:** The building blocks used in this study that have been solved by X-ray crystallography are listed. Although solved by crystallography, DHR5 was not not included in the set because homo-oligomerization was detected in the original study.

| |
|---|
| **DHR spacers:**<br>5CWB (DHR4), 5CWD (DHR7), 5CWF (DHR8), 5CWG (DHR10), 5CWH (DHR14),<br>5CWI (DHR18), 5CWJ (DHR49), 5CWK (DHR53), 5CWL (DHR54), 5CWM (DHR64),<br>5CWN (DHR71), 5CWO (DHR76), 5CWP (DHR79) and 5CWQ (DHR81). |
| **C2 homo-dimers:**<br>5KBA (Ank1C2), 5HRY (Ank3C2_1), 5J73 (2L4HC2_9), 5J0K (2L4HC2_23),<br>5J10 (2L4HC2_24) |

**Supplementary Table 5** - Expected oligomer masses versus those determined by native-MS. Differences between expected and measured values are within the limits of method accuracy and can be explained by a combination of adducts, oligomer size, signal quality, mass resolution and data processing settings. Artificial dimerization between oligomers can occur dependent on concentration and droplet size during the electrospray process. This was notably observed for designs D2-21.22 and D3-19.24.

| Design ID | Oligomeric state | Oligomer mass (expected, kDa) | Oligomer mass (native-MS, kDa) | Error (%) | Intensity (%) |
|---|---|---|---|---|---|
| D2-1.1B | 4 | 154.8 | 155.0 | 0.13 | 100 |
| D2-1.1D | 4 | 167.2 | 167.3 | 0.09 | 100 |
| D2-1.4H | 4 | 215.8 | 216.1 | 0.14 | 100 |
| D3-1.5A2 | 6 | 296.4 | 296.6 | 0.07 | 100 |
| D3-1.5C | 6 | 250.1 | 250.3 | 0.06 | 100 |
| D2-21.8 | 4 | 140.2 | 140.6 | 0.30 | 100 |
| D2-21.22 | 4 | 156.9 | 157.4 | 0.34 | 100 |
| | 8 (artificial 4-mer dimerization) | 313.8 | 315.0 | 0.39 | 10 |
| D2-21.26 | 4 | 160.0 | 160.7 | 0.37 | 100 |
| D2-21.29 | 4 | 147.2 | 147.7 | 0.35 | 100 |
| D3-19.14 | 6 | 209.1 | 209.9 | 0.40 | 100 |
| D3-19.19 | 6 | 208.4 | 209.2 | 0.39 | 100 |
| D3-19.20 | 6 | 188.8 | 189.3 | 0.53 | 100 |

| | | | | | |
|---|---|---|---|---|---|
| D3-19.24 | 6 | 182.5 | 183.4 | 0.49 | 100 |
| | 12 (artificial 6-mer dimerization) | 365.0 | 367.3 | 0.64 | 10 |

**Supplementary Text File 1:** A simple design script and command-line example applies symmetry and designs sidechains with whatever score-function is in beta at the time of use (beta_nov16 during this work). Symmetry definition files are provided in Supplementary Text Files S2 and S3.

```
<ROSETTASCRIPTS>
	<SCOREFXNS>
		<ScoreFunction name="sfx_hard_symm" weights="beta.wts" symmetric="1"  >
		</ScoreFunction>
	</SCOREFXNS>
	<TASKOPERATIONS>
		<InitializeFromCommandline name="init" />
		<RestrictIdentities name="nomutate_VIRTUAL" identities="XXX"
prevent_repacking="1" />
		<DisallowIfNonnative name="disallow_nonnative" disallow_aas="CPM" />
		<ReadResfile name="resfile_designable" filename="%%resfile%%" />
	</TASKOPERATIONS>
	<MOVERS>
	<SetupForSymmetry name="symmetry_setup"
definition="%%symdef%%"></SetupForSymmetry>
		<SymPackRotamersMover name="design_rotamers_resfile"
scorefxn="sfx_hard_symm"
task_operations="init,nomutate_VIRTUAL,resfile_designable,disallow_nonnative"></SymPackR
otamersMover>
	</MOVERS>
	<PROTOCOLS>
	<Add mover_name="symmetry_setup" />
		<Add mover_name="design_rotamers_resfile" />
	</PROTOCOLS>
</ROSETTASCRIPTS>
```

**bash$: <rosetta_scripts_path>** -ignore_zero_occupancy false -database **<rosetta_database_path>** -linmem_ig 10 -lazy_ig true -parser:protocol **<rosettascripts_xml_path>** -s **<pdb_path>** -native **<pdb_path>** -nstruct 1 -parser:script_vars symdef=**<D2_or_D3_symmetry_definition>** resfile=**<resfile>** -ex1 -ex2 -unmute all -out:pdb_gz true -out:path:all ./ -beta -overwrite -scorefile **<scorefile_name>**.sc

**Supplementary Text File 2:** A symmetry definition file for D2 symmetry, for redesign with Rosetta and RosettaScripts

```
symmetry_name d2
subunits 4
number_of_interfaces  3
E = 4*VRT0001 + 2*(VRT0001:VRT0002) + 2*(VRT0001:VRT0003) + 2*(VRT0001:VRT0004)
anchor_residue COM
virtual_transforms_start
start -1,0,0 0,1,0 0,0,0
rot Rz_angle 180.0
rot Rx_angle 180.0
rot Rz_angle 180.0
virtual_transforms_stop
connect_virtual JUMP1 VRT0001 VRT0002
connect_virtual JUMP2 VRT0002 VRT0003
connect_virtual JUMP3 VRT0003 VRT0004
set_dof BASEJUMP x(50) angle_x(0:360) angle_y(0:360) angle_z(0:360)
set_dof JUMP2 z(50) angle_z(0:90.0)
```

**Supplementary Text File 3**: A symmetry definition file for D3 symmetry, for redesign with Rosetta and RosettaScripts.

```
symmetry_name d3
subunits 6
number_of_interfaces  4
E = 6*VRT0001 + 6*(VRT0001:VRT0002) + 3*(VRT0001:VRT0004) + 3*(VRT0001:VRT0005) +
3*(VRT0001:VRT0006)
anchor_residue COM
virtual_transforms_start
start -1,0,0 0,1,0 0,0,0
rot Rz_angle 120.0
rot Rz_angle 120.0
rot Rx_angle 180.0
rot Rz_angle 120.0
rot Rz_angle 120.0
virtual_transforms_stop
connect_virtual JUMP1 VRT0001 VRT0002
connect_virtual JUMP2 VRT0002 VRT0003
connect_virtual JUMP3 VRT0003 VRT0004
connect_virtual JUMP4 VRT0004 VRT0005
connect_virtual JUMP5 VRT0005 VRT0006
set_dof BASEJUMP x(50) angle_x(0:360) angle_y(0:360) angle_z(0:360)
set_dof JUMP3 z(50) angle_z(0:60.0)
```
